# Ancestral secretory programs underlie the evolution of morphological innovations across Spiralia

**DOI:** 10.64898/2026.03.09.710468

**Authors:** Yitian Bai, Biyang Hu, Kunyin Jiang, Hong Yu, Lingfeng Kong, Shaojun Du, Shikai Liu, Qi Li

**Affiliations:** Key Laboratory of Mariculture, Ministry of Education, Ocean University of China, Qingdao 266003, China; Institute of Marine and Environmental Technology, Department of Biochemistry and Molecular Biology, University of Maryland School of Medicine, Baltimore, MD, USA; Qingdao Institute of Blue Seed Industry, Qingdao 266073, China

## Abstract

Understanding how morphological innovations arise from ancestral genetic and cellular systems remains a major challenge in evolutionary biology. Molluscan shells represent one of the most diverse morphological novelties within Spiralia and have been central to the ecological and evolutionary diversification of molluscs. However, the cellular basis of shell formation and the evolutionary origin of shell-forming cell types remain poorly understood. Here, we present a single-cell transcriptomic atlas of the Pacific oyster (*Crassostrea gigas*) mantle and show that molluscan shell-forming cells evolved through diversification built upon ancestral epithelial-secretory systems in spiralians. We identify six spatially segregated shell-forming epithelial cell types, and demonstrate that larval and adult shell-forming cells are developmentally independent and characterized by transcriptomes enriched for evolutionarily young genes and lineage-specific gene duplication. Cross-species cell-type comparisons further reveal that oyster shell-forming cells share transcriptional and functional features with epithelial-secretory programs across Spiralia. Together, we propose that a genetic foundation for shell formation was already present in the last common ancestor of Spiralia, and that molluscan shell diversity arose through repeated co-option of ancestral cellular programs coupled with novel genes. Our study provides a framework for understanding how ancient cellular architectures can be repeatedly reconfigured to generate morphological novelty during evolution.

## Introduction

Morphological innovations are lineage-restricted traits that occur in specific taxa but are absent in outgroups, and they represent a major driver of evolutionary diversification and ecological expansion (Erwin 2021). Despite their evolutionary importance, the mechanisms by which novel structures arise from pre-existing genetic and developmental systems remain poorly understood (Moczek 2008; Wagner and Lynch 2010; Carscadden et al. 2023). While genetic mutations and gene duplication provide raw material for evolutionary change, a central question concerns how such variation is reorganized within cellular systems to produce new functions and, ultimately, novel morphological traits (Xia et al. 2025). Therefore, identifying the cellular substrates that mediate the emergence of evolutionary novelty is critical for linking genetic variation to phenotypic innovation (Arendt et al. 2016; Tanay and Sebé-Pedrós 2021; Parker and Pennell 2025).

Spiralia is one of the most diverse bilaterian clades, encompassing molluscs, annelids, flatworms, and brachiopods, and related phyla that exhibit remarkable diversity in body plans, developmental strategies, and biomineralized structures (Laumer et al. 2015; Piovani and Marlétaz 2023; Sleight 2023). A striking feature of spiralian evolution is the repeated emergence of hardened extracellular structures, such as molluscan carbonate shells, annelid chaetae and tubes, and brachiopod shells (Wernstrom et al. 2022; Bai et al. 2026). Although these structures perform similar protective and structural roles, they differ markedly in composition, developmental origin, and morphology. This raised the question of whether they share ancestral cellular and transcriptional programs or instead evolved independently through convergent recruitment of distinct molecular mechanisms. The extensive diversity and recurrent evolution of these systems also provide an ideal framework for investigating the cellular and molecular bases of morphological novelty.

Molluscan shell represents a prominent example of evolutionary innovation and has contributed substantially to the ecological success and morphological diversification of molluscs (Wanninger and Wollesen 2019; Chen et al. 2025; Bai et al. 2026). Shell architectures are remarkably diverse across the phylum, ranging from the single shells in gastropods and tubular shells of scaphopods to paired valves in bivalves, whereas aculiferan molluscs bear diverse calcareous spicules, sclerites, or serial dorsal shell plates (Wanninger and Wollesen 2019). Comparative morphological and phylogenetic evidence suggested that conchiferans evolved from a uni-shelled ancestor, with the paired valves of bivalves representing a derived shell architecture within Mollusca (Wanninger and Wollesen 2019; Chen et al. 2025). Shell formation is orchestrated by the mantle, a specialized secretory epithelium that deposits mineral and organic components through highly coordinated cellular processes (Clark et al. 2020). Increasing evidences have indicated that this process is governed by a conserved regulatory architecture coupled with rapidly evolving effector genes, a combination that supports extensive morphological diversity in shell structure and composition (Aguilera et al. 2017; Zhao et al. 2018; Yarra et al. 2021; Bai et al. 2026). However, the cellular origin and evolutionary history of shell-forming cell types remain unclear. In particular, it is unknown whether molluscan shell-secreting cells represent lineage-specific cellular inventions or derive from ancestral cellular systems shared across Spiralia.

Recent advances in single-cell transcriptomics have enabled the classification of cell types beyond traditional histological models and quantitative comparisons of cell-type repertoires across species (Zhong et al. 2025), which provide unprecedented resolution for dissecting the cellular basis of novel phenotypes across animal evolution (Li et al. 2025; Piovani et al. 2025; Tommasini et al. 2025; Xiong et al. 2025). These methods offer a powerful framework for understanding the gene expression programs and cellular organization underlying morphological novelties. Here, we investigate single-cell transcriptomics, developmental comparisons, phylostratigraphy, and cross-species cell-type alignment to investigate the cellular and evolutionary origins of molluscan shell formation in the Pacific oyster (*Crassostrea gigas*). We construct a comprehensive single-cell atlas of the adult mantle, identify spatially distinct shell-forming epithelial cell populations, and compare them with secretory cell types across representative spiralians. By integrating gene-expression profiles with gene-age analyses, gene-duplication histories, and cross-species cell-type comparisons, we reveal that molluscan shell-forming cells represent lineage-specific innovations built upon ancestral epithelial and secretory programs of Spiralia, and extensively remodeled through lineage-specific molecular innovation. Our study supports a model in which morphological innovations arise through repeated co-option and diversification of ancestral cellular architectures, coupled with the recruitment of lineage-specific and duplicated genes.

## Results

### A single-cell atlas of the oyster mantle reveals spatially distinct shell-forming cell types

To explore the cellular and molecular characteristics underlying molluscan shell formation, we performed single-cell RNA sequencing (scRNA-seq) on the mantle of an adult *C. gigas*. After stringent quality control and doublet removal, we obtained transcriptomes from 37,411 cells and identified 33 cell clusters by applying the uniform manifold approximation and projection (UMAP) method for the dimensionality reduction (Figure 1A; Supplementary file 1). Cell populations were annotated by integrating differentially expressed gene markers with experimental evidence in oysters from published studies, with shell-forming cell types further validated by fluorescence *in situ* hybridization (FISH) in this study (Figure 1B-D; Supplementary file 2). We identified major mantle cell types, including shell-secreting epithelial cells (SECs), hemocytes, myocytes, ciliated cells, neurons, vesicular connective tissue cells, and proliferative cells, whereas clusters lacking characterized markers were retained as unknown populations. Among them, myocytes expressed multiple components of the contractile apparatus, including *CgSmhc*, *CgMELC*, *CgPM*, and *Tnnt*. Functional studies have shown that *CgSmhc* and *CgMELC* are required for muscle organization and myogenesis in oysters (Li et al. 2021a, b), whereas *CgPM*, encoding paramyosin, is a major structural component of invertebrate muscle thick filaments and is associated with oyster myogenesis (Li et al. 2021c). Ciliated cells expressed genes encoding ciliary and axonemal components, including *Tekt1*, *Crocc*, *Rsh4a-2*, and *Tbb-4*, whereas neuronal populations were characterized by neuropeptide-and neuroendocrine-associated genes such as *Myomodulin* and *7B2* (Piovani et al. 2023). Vesicular connective tissue cells expressed *STBD1*, a glycogen cargo receptor experimentally shown to mediate glycophagy and glycogen mobilization in oysters (Ren et al. 2026). Proliferative cells showed expression of the DNA-replication factor *PCNA*, previously validated as a marker of proliferating cells in oysters (Yu et al. 2022). Hemocytes were annotated using experimentally validated morphotype-associated markers, including *XBOX/TGC* for small granule cells, *LEVAR* for hyalinocytes, and *MOX* for vesicular cells (De La Forest Divonne et al. 2025).

**Figure 1.**
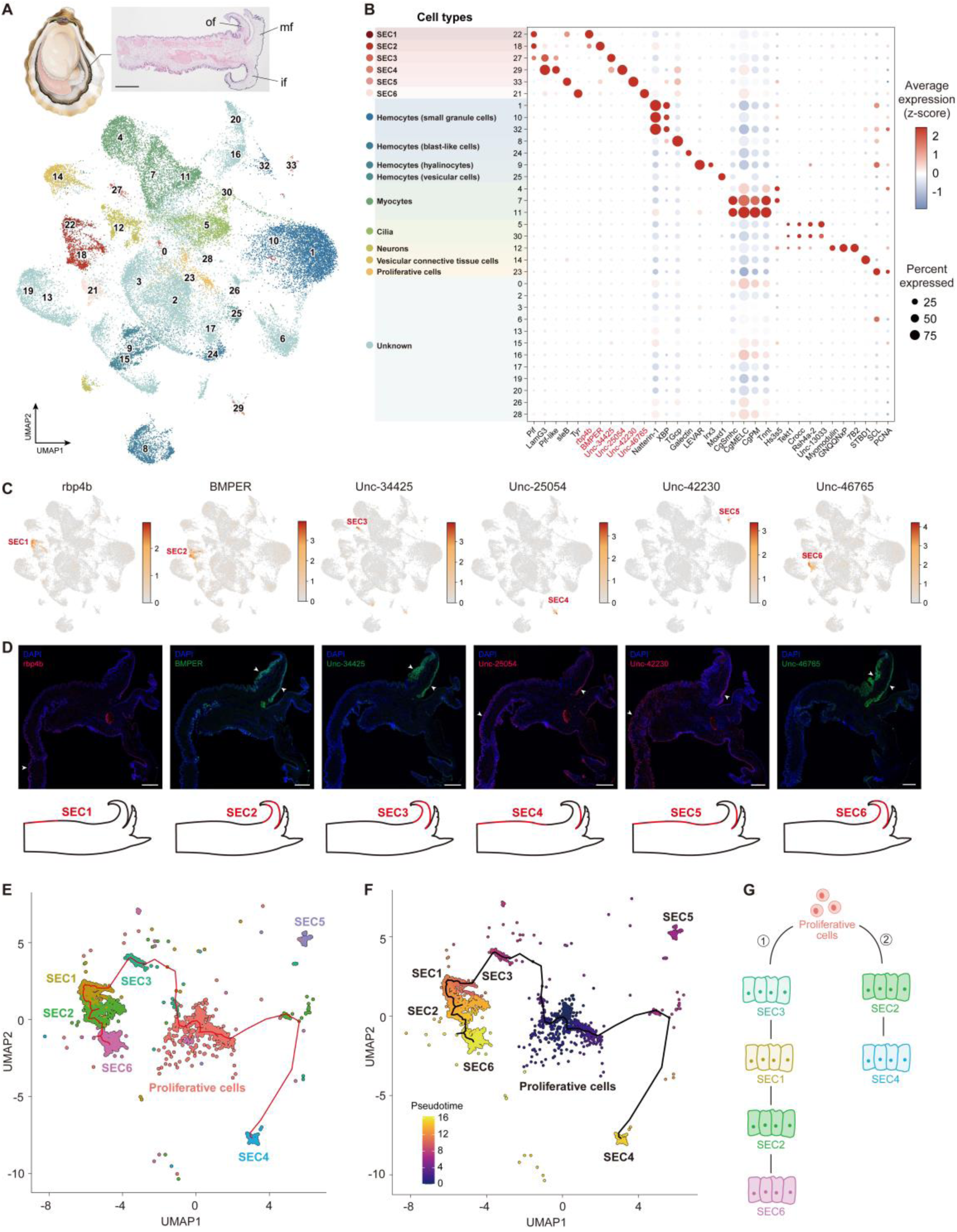
A single-cell atlas of the adult oyster mantle and spatial distribution of shell-forming cells. (A) UMAP visualization of mantle cells, with clusters colored according to their cell identities. A schematic illustration of oyster anatomy and H&E staining of a mantle cross-section are shown at the top. Abbreviations: of, outer fold; mf, middle fold; if, inner fold. Scale bar, 500 μm. (B) Dot plot showing the expression of representative marker genes across mantle cell clusters. SEC marker genes selected for FISH validation are highlighted in red. Markers used for annotation of other cell populations were compiled from published studies (Supplementary file 2). “Unc-” identifiers denote genes for which no sufficiently resolved standardized gene name was available. Clusters lacking sufficiently characterized marker genes are classified as “unknown”. (C) Feature plots showing the expression of six representative SEC marker genes used for spatial validation of SEC populations: *rbp4b* (SEC1), *BMPER* (SEC2), *Unc-34425* (SEC3), *Unc-25054* (SEC4), *Unc-42230* (SEC5), and *Unc-46765* (SEC6). (D) FISH of the six SEC marker genes shown in (C). Nuclei were counterstained with DAPI. Arrowheads indicate representative FISH signals. Schematic mantle cross-sections below each image summarize the spatial expression domain of the corresponding marker and the corresponding SEC population. Scale bars, 250 μm. (E) UMAP visualization of SEC and proliferative cell populations used for pseudotime analysis. Red lines indicate the inferred principal trajectories. (F) Pseudotime trajectories of SEC and proliferative cell populations, with proliferative cells specified as the root. Colors indicate pseudotime values, and black lines indicate the inferred trajectories. (G) Schematic summary of the major pseudotime-inferred transcriptional trajectories connecting proliferative cells and SEC populations. One branch connects proliferative cells with SEC3, SEC1, SEC2, and SEC6, whereas a second branch extends toward SEC4. SEC5 was not included because it was disconnected from the major inferred trajectories.

The overall identity of SECs was assigned based on the spatial expression of previously characterized marker genes associated with oyster shell formation, such as *Pif*, *Pif-like*, *LamG3*, *sleB*, and *Tyr*, which are involved in shell matrix secretion and biomineralization (Zhu et al. 2021; Bai et al. 2023). Within the SECs, six transcriptionally distinct SEC populations were resolved, corresponding to clusters 22, 18, 27, 29, 33, and 21 (SEC1–SEC6, respectively). To independently validate these populations and determine their spatial distributions within the mantle, we selected representative marker genes for each SEC population: *rbp4b* for SEC1, *BMPER* for SEC2, *Unc-34425* for SEC3, *Unc-25054* for SEC4, *Unc-42230* for SEC5, and *Unc-46765* for SEC6 (Figure 1B-C; Supplementary file 2). FISH revealed distinct spatial expression domains for these markers along the mantle epithelium and outer fold (Figure 1D). Among them, SEC1 was predominantly distributed along the outer epithelium of the pallial mantle. In contrast, SEC2, SEC3, and SEC6 were enriched toward the epithelium of outer fold, including the periostracal groove, which is involved in periostracum formation. SEC4 and SEC5 showed broad expression along the pallial outer epithelium and on the inner surface of the outer fold. Dual-color FISH further confirmed that the transcriptionally related SEC1 and SEC2 populations exhibited distinct spatial localization, whereas the SEC3 and SEC4 markers showed spatially differential distributions with only limited overlap (Figure 1-figure supplement 1). These results provide the support for the subdivision of the shell-secreting epithelium into regionally specialized SEC populations and reveals pronounced molecular and spatial heterogeneity within the adult mantle. These spatially regionalized expression programs likely reflect functional compartmentalization for biomineralization process, corresponding to the secretion of distinct organic matrices and the formation of different shell layers and microstructures (Marie et al. 2012; Shimizu et al. 2022; Bai et al. 2023).

To explore transcriptional relationships among shell-forming cell types, we performed a pseudotime trajectory analysis on the six SEC populations together with proliferative cells (Figure 1E-F). Proliferative cells were specified as the root based on their strong expression of cell-cycle-associated genes, such as *PCNA* (Figure 1B). The inferred principal graph revealed two major transcriptional branches connecting proliferative cells with SEC populations. One branch extended through SEC3 and subsequently connected SEC1, SEC2, and SEC6, whereas a second branch extended toward SEC4 (Figure 1G). SEC5 formed a transcriptionally distinct population that was disconnected from the principal trajectory and was therefore not assigned to either branch. These results reveal a branched transcriptional organization among SEC populations, linking their spatial segregation within the mantle to distinct transcriptional states that may reflect regional specialization in shell formation.

### Larval and adult shell-forming cells deploy distinct and rapidly evolving molecular programs

During molluscan development, larval and adult shells exhibit striking differences in shapes, microstructures and shell matrix protein (SMP) repertoires (McDougall and Degnan 2018; Zhao et al. 2018; Cavallo et al. 2022). We thus reasoned whether shell-forming cells at the larval and adult stages retain similar transcriptional identities. To characterize larval shell-forming cells, we re-analyzed the publicly available scRNA-seq data of *C. gigas* trochophore (Piovani et al. 2023), and obtained a single-cell atlas comprising 6,978 larval cells (Figure 2A; Supplementary file 3). Within this dataset, four larval shell gland populations, together with proliferative cells, neurons, hemocytes, ciliated cells, myocytes, were identified based on their transcriptional profiles and the expression of marker genes (Figure 2B; Figure 2-figure supplement 1; Supplementary file 2). We further analyzed the expression patterns of oyster SMP genes at both bulk (Zhang et al. 2012; Lian et al. 2025) and single-cell levels (Figure 2-figure supplement 2). Larval and adult SMP gene sets exhibited markedly different temporal and tissue-expression profiles across oyster development (Figure 2-figure supplement 2A, D). Larval SMP genes were predominantly expressed in trochophore shell gland cells but were barely detected in SECs (Figure 2-figure supplement 2B-C). Conversely, adult SMP genes were preferentially expressed in adult SECs and showed limited expression in larval shell gland cells (Figure 2-figure supplement 2E-F). These reciprocal expression biases support extensive stage-specific deployment of biomineralization effector genes during larval and adult shell formation (Zhao et al. 2018; Bai et al. 2026).

**Figure 2.**
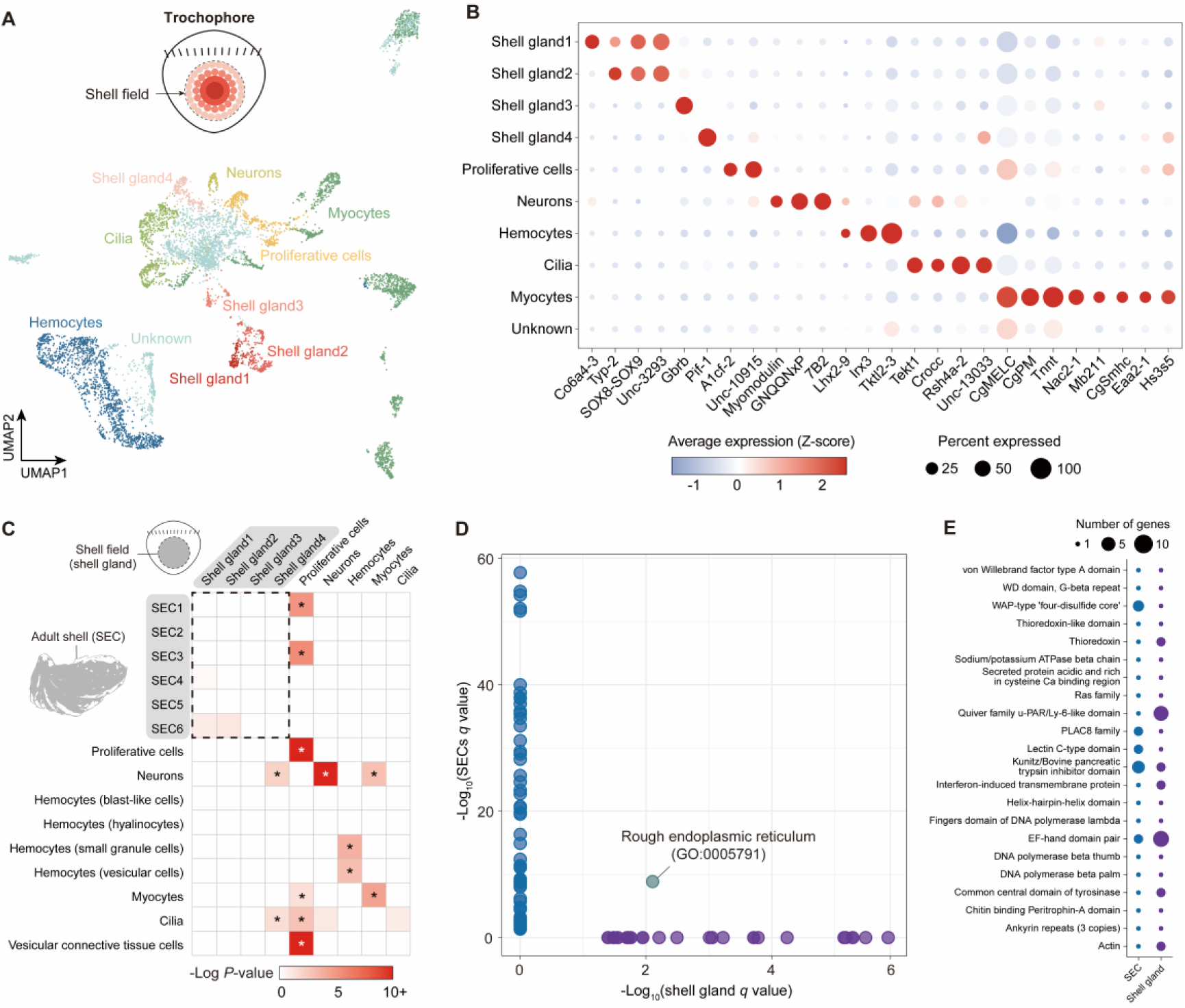
Distinct shell-forming cell types between larval and adult stages in *C. gigas*. (A) UMAP visualization of integrated single-cell transcriptome of the trochophore. The schematic at the upper left illustrates the position of the shell field and the distribution of shell gland cell populations in the trochophore dorsal side. (B) Dot plot showing the expression of representative marker genes of the major trochophore cell populations. Detailed information of marker genes used for cell-type annotation is provided in Supplementary file 2. Clusters lacking sufficiently characterized marker genes are classified as “unknown”. (C) Pairwise comparison of marker-gene overlap of larval and adult (mantle) cell types. For each pair, the box is shaded by statistical significance (-log10 scale of *P* values by hypergeometric test). Asterisks indicate significant overlap (*P* < 0.05). The dashed box highlights comparisons between larval and adult shell-forming cell types. (D) Comparative GO enrichment analysis of marker genes from adult SECs and larval shell gland cells. Each point represents a GO term significantly enriched in at least one of the two groups (*q* < 0.05, FDR-adjusted). GO terms significantly enriched only in SECs are shown in blue, those enriched only in shell gland cells in purple, and terms significantly enriched in both groups in green. Rough endoplasmic reticulum (GO:0005791) was the only GO term significantly enriched in both larval shell gland cells and adult SECs. Detailed information of GO enrichment is provided in Supplementary file 5. (E) Comparison of shared protein domains identified among proteins encoded by shell gland and SEC marker genes. Dot size indicates the number of genes encoding proteins containing each domain. Abbreviations: SEC, shell-secreting epithelial cells.

Comparative analysis further demonstrated minimal overlap between marker genes specifically enriched in larval shell glands and those enriched in adult SECs, with no significant correlation between their expression profiles across cell types (Figures 2C; Figure 2-figure supplement 3; Supplementary file 4). In contrast, significant marker-gene overlaps were observed between several corresponding non-shell-forming cell types in the trochophore and adult mantle, including proliferative cells, neurons, hemocytes, and myocytes, indicating that these cell types retain relatively conserved transcriptional identities between the two datasets. Moreover, GO enrichment analysis of marker genes revealed largely distinct functional profiles between larval shell glands and adult SECs, with rough endoplasmic reticulum representing the only shared GO term (Figure 2D; Supplementary file 5). These findings indicate substantial transcriptional divergence between larval and adult shell-forming systems, with the two shells secreted by developmentally independent cell populations that deploy largely stage-specific gene repertoires. Nevertheless, comparison of protein-domain composition revealed several domains shared between shell gland-and SEC-enriched proteins, including von Willebrand factor type A (VWA), WAP-type four-disulfide core, EF-hand, Kunitz/BPTI trypsin inhibitor, chitin-binding Peritrophin-A (CB), and tyrosinase domains (Figure 2E; Supplementary file 6). Many of these domains have been proposed as components of conserved biomineralization toolkits (Aguilera et al. 2017; Yarra et al. 2021; Bai et al. 2023), whereas VWA and CB domains have been identified in both larval and adult SMP repertoires (Zhao et al. 2018). Thus, larval and adult shell-forming cells share partially overlapping protein-domain architectures associated with shell biomineralization, suggesting conservation of some biochemical modules despite extensive divergence at the gene level.

Phylostratigraphic analyses of single-cell transcriptomes indicated that both larval shell glands and adult SECs exhibited the younger transcriptome ages, defined as a relative enrichment of expressed genes with more recent evolutionary origins inferred from phylostratigraphy, compared to many other cell types in larvae and adult mantle (Figure 3A-B). Consistent with this pattern, enrichment analysis of cell-type marker genes across successive phylostrata revealed distinct contributions of lineage-restricted genes to both larval shell gland and adult SEC populations (Figure 3C-D). Notably, several shell gland populations and SECs showed significant enrichment for genes originating at relatively recent nodes within Mollusca, Bivalvia, and Ostreidae. These results suggest that both larval and adult shell-forming programs have independently incorporated substantial numbers of evolutionarily young genes and evolved independently, each recruiting distinct sets of novel genes into shell formation. However, by extending this approach to the bulk level, a peak of young gene expression was observed during the trochophore stage, whereas adult shells exhibited among the oldest transcriptome profiles and the digestive gland exhibited among the youngest ones (Figure 3-figure supplement 1). The apparent difference between the adult single-cell and bulk-tissue analyses likely reflects their different levels of resolution: single-cell analysis resolves transcriptionally specialized cell populations within the mantle, whereas bulk transcriptomes average expression across heterogeneous cellular compositions.

**Figure 3.**
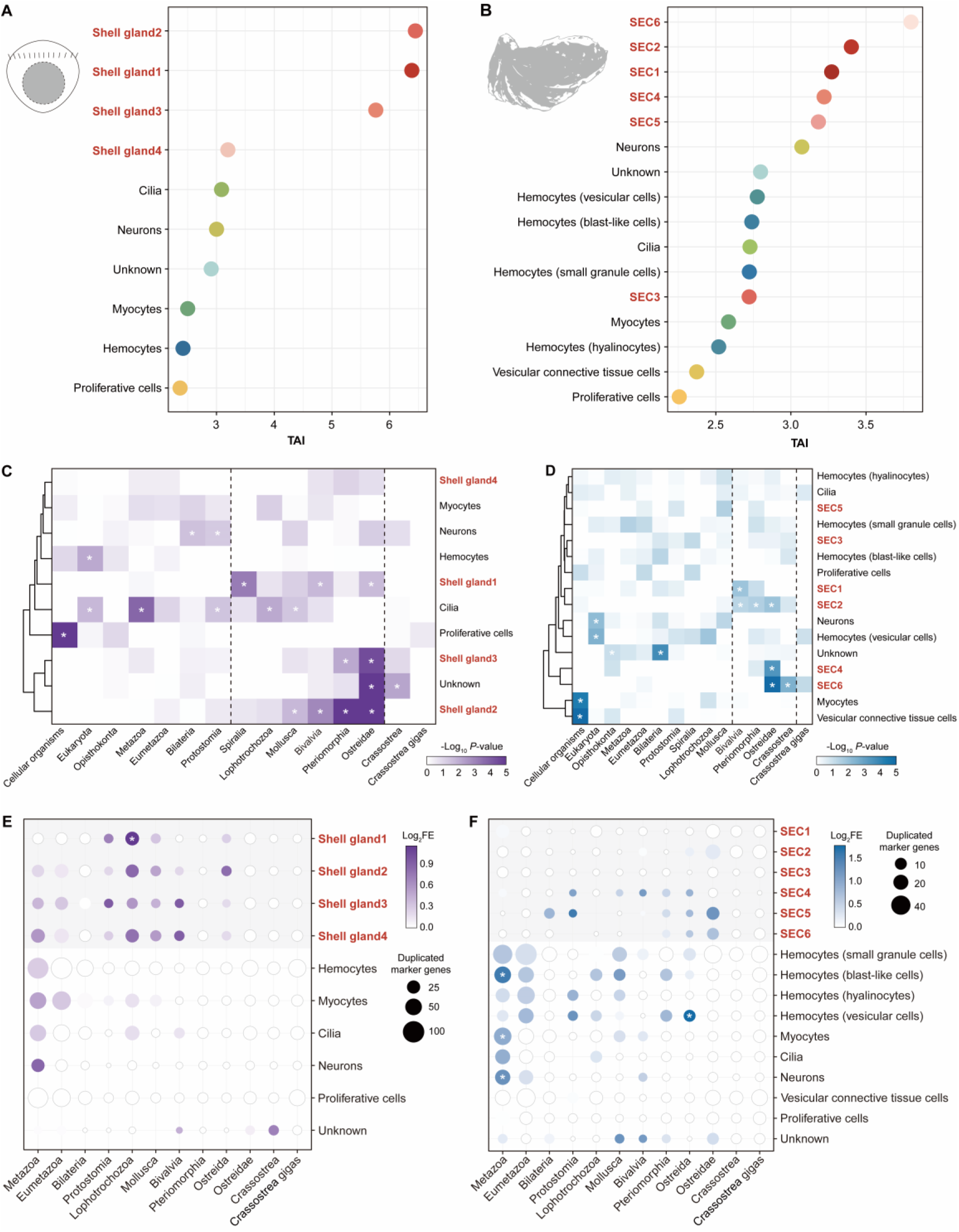
Gene age and duplication of larval and adult shell-forming cell types in *C. gigas*. (A, B) Transcriptome age index (TAI) of cell types in the trochophore (A) and adult mantle (B). Higher TAI values correspond to “younger” gene ages. Shell-forming cell types are marked in red. (C, D) Hypergeometric enrichment of genes that originated at successive phylostrata in the trochophore (C) and adult mantle (D) cell types. The dashed boxes highlight recently evolved genes of shell-forming cell types in the trochophore and adult mantle, respectively. Shell-forming cell types are marked in red. Asterisks indicate significant enrichment (BH-adjusted *P* < 0.05). (E, F) Phylogenetic distribution and enrichment of duplicated marker genes across inferred duplication nodes in trochophore (E) and adult mantle (F) cell types. Dot size represents the number of duplicated marker genes assigned to each duplication node, and color indicates the log2 fold-enrichment (log2FE) relative to the corresponding background gene set. Shell-forming cell types are marked in red, and gene duplication associated with these cell types are highlighted by grey shading. Asterisks indicate statistically significant enrichment (BH-adjusted *P* < 0.05).

To further assess the contribution of gene duplication to shell-forming cell diversification, we examined the phylogenetic distribution of duplication events among cell-type marker genes. Duplicated marker genes were detected across multiple evolutionary nodes in both larval shell gland and adult SEC populations, including lineage-specific events within Lophotrochozoa, Mollusca, Bivalvia, and Ostreidae (Figure 3E-F; Supplementary file 7). Shell gland1 showed significant enrichment for genes derived from duplications at the Lophotrochozoa node, whereas other shell gland and SEC populations exhibited more heterogeneous duplication patterns. Notably, duplicated genes expressed in shell-forming populations included several gene families associated with extracellular matrix secretion and biomineralization, including alkaline phosphatases, tyrosinases, chitin synthases and chitin-associated proteins, collagen-and VWA-containing proteins, as well as proteins containing WAP and Kunitz/BPTI domains (Supplementary file 7). GO enrichment analysis further revealed that duplicated genes in larval shell gland cells were mainly associated with epithelial junction organization and transmembrane transport, whereas those in adult SECs were enriched for translation and protein targeting to the endoplasmic reticulum and membrane (Supplementary file 8). These results suggest that lineage-specific gene duplication provides an additional source of molecular innovation that contributes to the functional diversification and rapid evolution of larval and adult shell-forming programs.

### Spiralian epithelial-secretory molecular programs underlie molluscan shell-forming cell innovation

To trace the evolutionary origins of molluscan shell-forming cells, we performed pairwise cross-species cell-type comparisons between *C. gigas* and four representative spiralians, including the annelid *Pristina leidyi*, the flatworms *Dugesia japonica* and *Schmidtea mediterranea*, and the chaetognath *Paraspadella gotoi* (Figures 4A-D, Figure 4-figure supplement 1). We found transcriptional similarities between several oyster SEC populations and secretory or epithelial-related cell populations in the other spiralians, although the alignment patterns differed among species. In *P. leidyi*, SEC populations showed affinities with chaetal-sac, epidermal, and eleocyte populations, whereas SECs in the two flatworms primarily aligned with distinct secretory cell populations. In *P. gotoi*, SECs showed transcriptional similarities primarily to epidermal and bristle-associated populations. These results indicate that molluscan shell-forming cells share transcriptional features with several secretory and epithelial cell populations across distant spiralian lineages.

**Figure 4.**
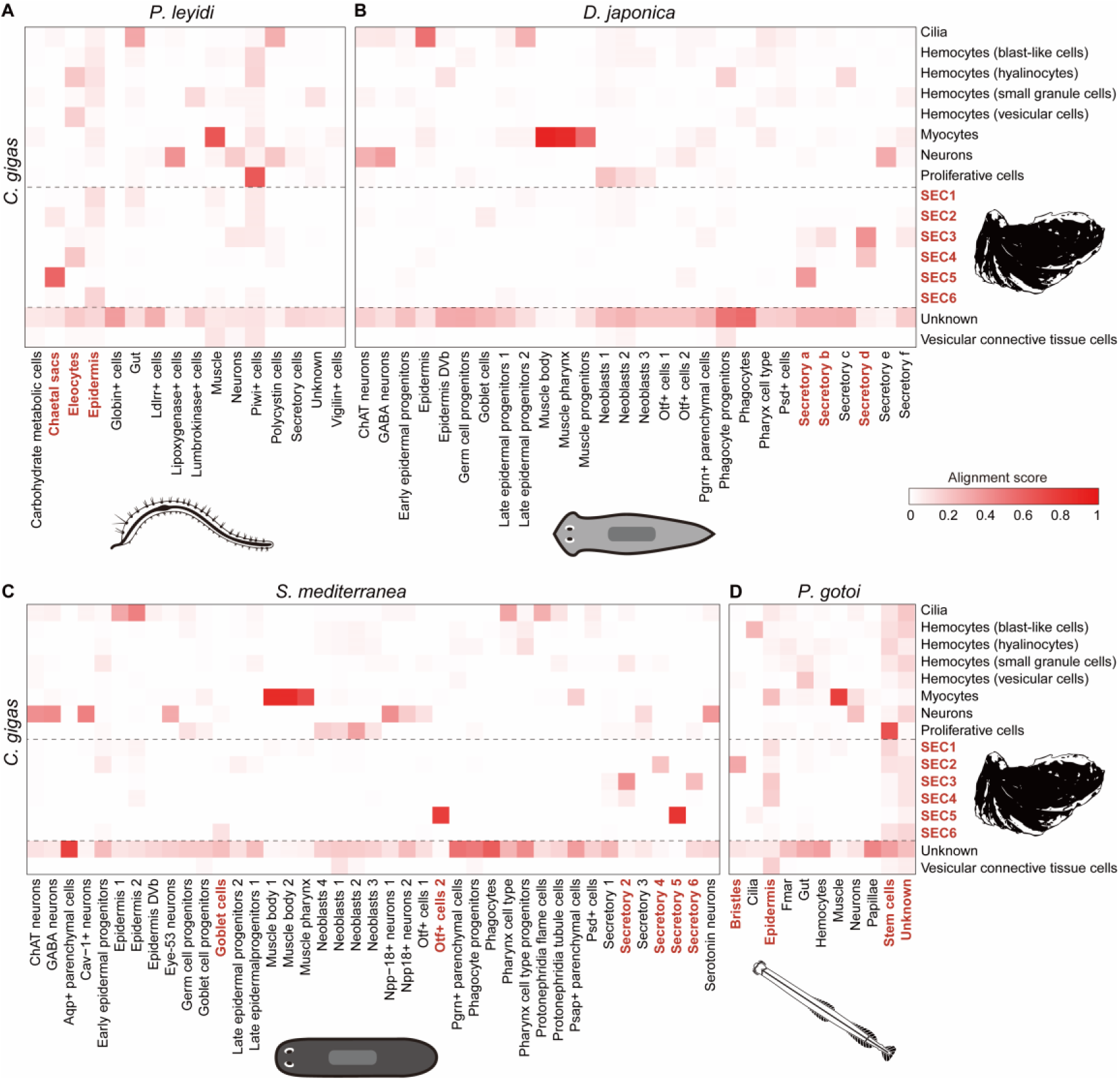
Cross-species cell-type alignment between adult *C. gigas* and other spiralians. (A-D) Pairwise SAMap alignments between *C. gigas* and (A) the annelid *P. leidyi*, (B) the flatworm *D. japonica*, (C) the flatworm *S. mediterranea*, and (D) the chaetognath *P. gotoi*. The scale bar represents the SAMap alignment score, defined as the average number of mutual nearest cross-species neighbors for each cell relative to the maximum possible number. Dashed boxes highlight the alignment with SECs of *C. gigas*.

To determine which genes contributed to these cross-species similarities, we investigated the homologous gene pairs contributing to the SAMap alignments (Supplementary file 9). SEC-associated alignments with *P. leidyi*, *D. japonica*, *S. mediterranea*, and *P. gotoi* involved 16, 9, 11, and 124 unique *C. gigas* genes, respectively (Figure 4-figure supplement 2A). Overlap among these gene sets was limited, with only two oyster genes represented in all four comparisons: *LOC105344059*, encoding a glioma pathogenesis-related protein 1 containing a CAP domain, and *LOC117683433*, encoding a biological adhesion-related protein. Other recurrent genes included *LOC105328359* (peptidase inhibitor), *LOC105328360* (glioma pathogenesis-related protein 1-like), and *LOC105344061* (CRISP-family protein), whereas most alignment-associated genes were restricted to a subset of species comparisons. Functional enrichment further indicated that the molecular basis of the alignments differed among species (Figure 4-figure supplement 2B). Gene-pair sets associated with the two flatworms were enriched for stress-response and vesicle/lysosomal membrane functions, whereas several terms related to translation, ribosomal functions, endoplasmic reticulum (ER) protein targeting, and RNA metabolism were enriched in those of *P. leidyi* and *P. gotoi*. Despite the limited conservation of individual gene identities, these recurrent molecular features involve processes related to secretion and epithelial activity. Thus, the cross-species similarities reflect partial conservation at the level of cellular function rather than strict conservation of the underlying gene repertoire. These findings suggest that ancestral functions associated with secretion and epithelial matrix production have been retained during the evolution of molluscan shell-forming cells, consistent with their secretory roles in biomineralization.

Notably, we found multiple cell types, including myocytes, neurons, and proliferative cells, aligned between oyster and other spiralian species (Figure 4). These conserved cell populations are likely to be homologous across Spiralia (Piovani et al. 2025), as supported by the shared expression patterns of homologous genes (Figure 5A; Supplementary file 10). Moreover, oyster SECs shared 18 gene pairs with chaetal-sac cells in *P. leidyi* and 7 and 13 gene pairs with secretory cells in *D. japonica* and *S. mediterranea*, respectively. Importantly, extensive gene-pair support was observed in *P. gotoi*, where SECs shared 111 gene pairs with bristle-associated cells and 149 with epidermal cells (Figure 5A; Supplementary file 10). Compared with the highly conserved muscle, neuronal and proliferative populations, these variable levels of gene-pair support suggest that SEC-related similarities are not based on a uniformly conserved cell-type transcriptome, but are instead consistent with partial retention or repeated co-option of ancestral epithelial-secretory molecular features across spiralian lineages. Moreover, Lineage-specific bursts of gene novelty were detected in SECs, particularly along the Bivalvia, Pteriomorphia, Ostreidae, and *Crassostrea* lineages (Figure 3D), suggesting the stepwise integration of lineage-specific novelties of effector genes into shell formation. Together with the recurrent secretory and epithelial functions (Figure 4-figure supplement 2), these results support a model in which deeply conserved cellular systems, such as muscle, neural, and stem cell types, were already present in the spiralian ancestor, whereas lineage-specific secretory and epithelial specializations subsequently diversified through modification and redeployment of ancient molecular functions (Figure 5B). Under this model, molluscan shell-forming cells represent a derived cellular specialization built upon an ancestral epithelial-secretory framework, with progressive incorporation of lineage-specific effector genes contributing to the diversification of shell-forming cell types and the complexity of molluscan shells.

**Figure 5.**
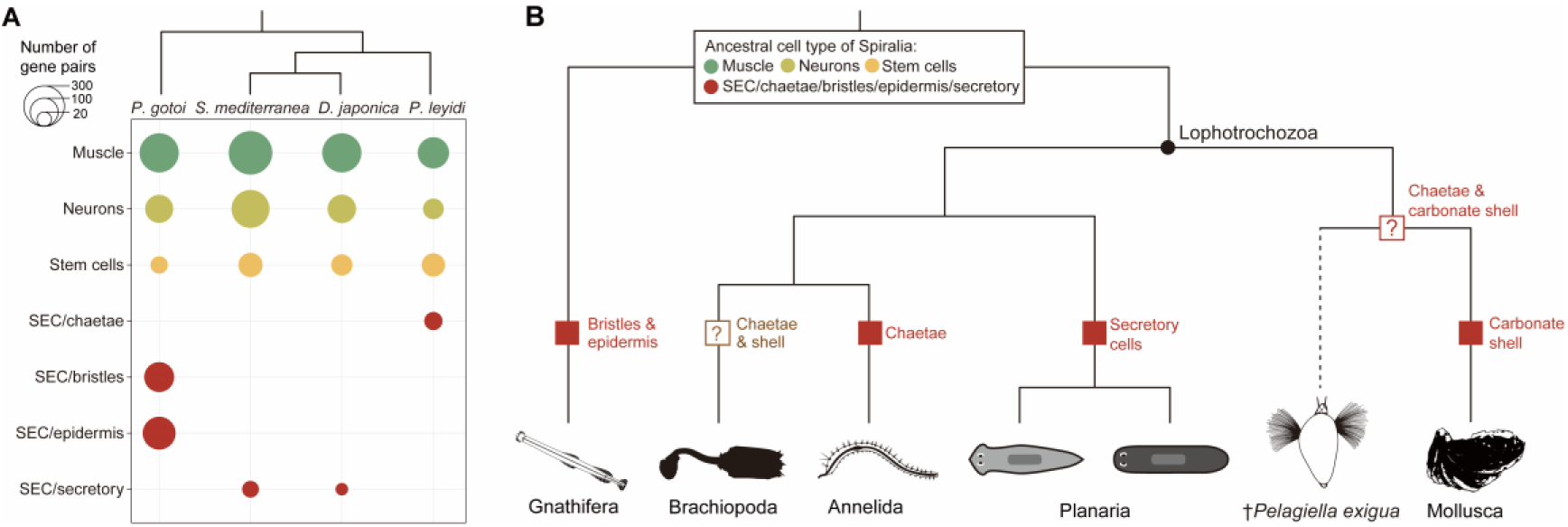
Cellular systems and evolutionary diversification underpin spiralian morphological innovations. (A) Cross-species comparison of homologous gene pairs supporting SAMap alignments between broad cell-type categories in adult *C. gigas* and four representative spiralians: the chaetognath *P. gotoi*, the flatworms *S. mediterranea* and *D. japonica*, and the annelid *P. leidyi*. Circle size represents the number of homologous gene pairs associated with each cell-type alignment. Muscle, neuronal, and stem/proliferative populations show broad cross-species correspondence, whereas SEC-related alignments involve chaetal-sac cells in *P. leidyi*, secretory cells in the two flatworms, and bristle-associated or epidermal populations in *P. gotoi*. Complete gene-pair information is provided in Supplementary file 10. (B) Schematic evolutionary model illustrating the proposed relationships between deeply conserved cellular systems and lineage-specific epithelial-secretory specializations across Spiralia. Muscle, neuronal, and stem/progenitor cell types are depicted as broadly conserved ancestral cellular systems, whereas red symbols indicate derived epithelial or secretory populations associated with hardened structures or specialized secretion in different spiralian lineages, including chaetognath bristle-associated and epidermal cells, annelid chaetal cells, planarian secretory cells, and molluscan shell-secreting epithelial cells. Question marks indicate two unresolved evolutionary problems in spiralian characters: one concerns the brachiopods, where the homology between their shells and molluscan carbonate shells, as well as the relationship between brachiopod and other spiralian chaetae, remain unclear; the other relates to the Cambrian fossil *Pelagiella exigua*, a potential stem-group mollusc that possibly possessed both chaetae and a carbonate shell, leaving open whether it represents the ancestral molluscan morphology or an independent lineage that evolved both traits convergently.

## Discussion

Spiralia is an extremely species-rich clade that diverged from the other major protostome lineage Ecdysozoa during the early Cambrian period, approximately 550 million years ago (Peterson et al. 2004). Since then, spiralians have undergone remarkable diversification of body plans, developmental modes, and biomineralized structures. Many of these morphological novelties are characterized by lineage-specific hardened structures, including molluscan carbonate shells, annelid chaetae and chaetognath bristles, and brachiopod shells (Figure 5B), raising the question of whether they arose through the invention of entirely new cellular systems or through modification of pre-existing cellular and molecular architectures (Piovani and Marlétaz 2023; Sleight 2023). By resolving the cell-type composition responsible for oyster shell formation and integrating developmental, evolutionary and cross-species comparative analyses, our study reveals that molluscan shell formation represents derived cellular specializations assembled through the diversification of ancestral epithelial-secretory functions across Spiralia, together with extensive lineage-specific molecular innovation. Several SEC types (SEC1, SEC2, SEC4 and SEC6) exhibit relatively weak cross-species alignments and deploy particularly young transcriptomes, whereas SEC3 and SEC5 show strong transcriptional affinities to secretory or epithelial cell types in other spiralians (Figure 3B and 4). This observation is consistent with the viewpoint that cellular novelties often emerge through the recombination or redeployment of conserved gene expression programs, while subsequently integrating rapidly evolving effector repertoires (Parker and Pennell 2025). The transcriptional trajectories inferred that SEC6 served as a terminally branched population associated with rapidly evolving secretomes (Figure 1E-G), further supporting that mantle shell-forming cells comprise a mosaic of ancestrally related secretory cell types and more recently derived effector cell types that have diversified within molluscs (Jackson et al. 2006; Kocot et al. 2016). Distinct cellular zones within the mantle epithelium are dedicated to secreting specific organic matrices and controlling mineral polymorphs through differential gene expression (Herlitze et al. 2018; Sleight et al. 2020; Shimizu et al. 2022; Bai et al. 2023). The pronounced compartmentalization of SECs along the mantle epithelium suggests that region-specific modification of an ancestrally homogeneous epithelium enabled the emergence of distinct biomineralizing modules that pattern shell architecture.

Similarly to adult mantle, the larval shell gland also exhibits a modular organization comprising distinct cell populations (Liu et al. 2020; Piovani et al. 2023). Molluscan trochophore larvae and mantle show transcriptomes enriched for evolutionarily young genes, underlying shell formation and diversity (Wu et al. 2019; Wang et al. 2020). This pattern is consistent with the rapid evolution of biomineralization effectors, many of which are lineage-restricted or rapidly evolving genes encoding secreted proteins (Jackson et al. 2006; Kocot et al. 2016; Aguilera et al. 2017). Such molecular turnover may facilitate the diversification of shell-forming cellular programs and contribute to lineage-specific shell architectures (Arivalagan et al. 2017; Cavallo et al. 2022). However, larval shell-gland cells and adult SECs show no significant transcriptional correlation, consistent with the previous report that larval and adult shells rely on largely distinct SMPs and gene repertoires (Zhao et al. 2018). Moreover, both larval and adult shell-forming systems process young transcriptomes enriched for lineage-specific genes. Together, these observations indicate that larval and adult shell-forming systems have undergone independent rounds of molecular innovation, repeatedly recruiting lineage-restricted effectors into stage-specific biomineralization programs (Bai et al. 2026). In addition, shared protein-domain architectures between the two stages indicates that extensive gene-level turnover can occur within a more constrained biochemical framework. This stage specificity does not necessarily imply complete molecular independence between larval and adult shell-forming systems. A recent study reported that a stem cell niche within the adult scallop shells expressed several larval SMP genes and are involved in shell formation, which were identified through sequence homology with oyster SMPs (Lian et al. 2025). These genes are also highly expressed during the umbo larva stages and subsequent developmental stages in the oyster and scallop (Lian et al. 2025), and may represent a subset of conserved proliferative or stem cell-related genes broadly shared among bivalves, rather than the complete suite of SMPs in scallop shells. It is likely that a large number of lineage-specific SMPs in scallops remain unidentified, exhibiting stage-specific expression patterns for larval and adult shell formation. Further in-depth studies across a broader range of molluscan species, integrating SMP identification by mass spectrometry and cell-type annotation with lineage-specific biomineralization genes, will be essential for resolving these lineage-and stage-specific biomineralization programs.

Gene duplication can generate lineage-specific paralogs from ancient gene families, allowing pre-existing molecular functions to be expanded, partitioned, or recruited into new cellular contexts. Such processes are widespread in spiralians and are particularly relevant to molluscan biomineralization (Piovani et al. 2025; Bai et al. 2026). Our results demonstrate that duplicated genes recruited into shell-forming populations include families associated with extracellular matrices, chitin-related functions, secretion, and biomineralization. Gene age and duplication age capture complementary dimensions of shell evolution, as a gene may belong to an ancient family yet represent a recently derived paralog with a lineage-specific cellular role. Thus, repeated expansion and diversification of biomineralization-related gene families may contribute to molecular novelty alongside *de novo* gene emergence, which contributed to shell diversity (Aguilera et al. 2017; Yarra et al. 2021; Bai et al. 2026). Together, *de novo* gene emergence and lineage-specific duplication provide complementary routes through which ancestral cellular systems can acquire derived biomineralization functions, allowing shell-forming cell states to evolve through progressive remodeling of ancestral cellular machinery rather than its wholesale replacement (Xia et al. 2025; Parker and Pennell 2025).

TAI analyses summarize the evolutionary ages of genes expressed in a given sample and are therefore inherently influenced by its cellular composition and transcriptional state (Domazet-Lošo et al. 2007; Drost et al. 2018; Wang et al. 2020). In this study, our adult single-cell dataset is restricted to the mantle, which limits our ability to place the evolutionary ages of mantle cell types within a whole-organism context. Within the mantle, neuronal and myocyte populations provide useful internal references because neural and muscle cell identities represent deeply conserved metazoan cell types (Arendt et al. 2016; Tarashansky et al. 2021), allowing the relatively young transcriptomic profiles of several SECs to be evaluated against more broadly conserved cellular identities. However, these mantle-associated neurons and myocytes cannot substitute for dedicated nervous or muscle tissues, which contain additional specialized cell populations and transcriptional states. This limitation is further illustrated by the difference between our single-cell and bulk-tissue analyses (Figure 3-figure supplement 1), as individual SEC populations are enriched for evolutionarily younger genes, whereas the adult shell shows an overall old transcriptome-age profile at the bulk level, emphasizing how tissue-level measurements average across heterogeneous cellular compositions. Thus, a whole-organism adult single-cell atlas, including dedicated nervous, muscular, digestive, and other organ systems, will ultimately be required to determine the evolutionary age of shell-forming cells relative to the complete oyster cell-type repertoire.

Secretory and epithelial functions are widespread across Spiralia, and several molecular systems involved in extracellular matrix production, secretion, chitin-associated processes, and biomineralization have been repeatedly recruited during the evolution of hardened structures (Sun et al. 2020; Sun et al. 2021; Wernstrom et al. 2022; Barrera Grijalba et al. 2025). Consistent with our cross-species comparisons, this pattern suggests that evolutionary innovation may proceed through modification and redeployment of ancestral cellular capabilities while allowing extensive turnover of the underlying gene repertoire. Such a framework is compatible with models of cell-type evolution in which conserved cellular functions provide a substrate for lineage-specific recruitment of novel genes, duplicated paralogs, and new regulatory interactions (Piovani and Marlétaz 2023; Sleight 2023; Parker and Pennell 2025). The diversification of SECs fits this paradigm, where ancestral epithelial-secretory functions were progressively modified through genetic innovation, giving rise to distinct biomineralizing cell populations with specialized function of shell formation. Thus, the spiralian last common ancestor likely possessed broadly conserved cellular systems, including muscle, neuronal, stem, and epithelial-secretory functions, that provided substrates for lineage-specific cellular diversification in each lineage (Figure 5B). Our findings suggested that morphological innovations arise through modifications of ancestral cell-type architectures and the integration of lineage-specific genes, thereby generating derived cellular functions rather than entirely novel cellular systems. Furthermore, these innovations may be governed by the presence and activation of regulatory elements that control gene expression (Martin-Zamora et al. 2023; Parey et al. 2024; Li et al. 2025; Piovani et al. 2025). Future integrative approaches such as single-cell ATAC-seq, by mapping open chromatin regions, and cis-regulatory elements at cellular resolution, will enable cross-species reconstruction of regulatory networks and provide more definitive evidence for cell type homology and conserved regulatory programs underlying evolutionary innovations.

More broadly, an unresolved question is whether brachiopod shells and chaetae share the common evolutionary origins with the molluscan carbonate shells and annelid chitinous chaetae (Figure 5B). Although convergent evolution of effector gene repertoires is well documented in biomineralization (Jackson et al. 2010; Luo et al. 2015; Murdock 2020; Sleight 2023), our results suggest that the spiralian hardened structures, including molluscan shells, annelid chaetae, and chaetognath bristles, may trace back to an ancestral secretory cell-type framework. Fossil and comparative evidences further indicated that brachiopod and molluscan skeletons repeatedly evolved through the co-option of organic, often chitinous, precursors rather than from a biomineralized common ancestor (Murdock 2020). Nonetheless, future single-cell transcriptomic studies of brachiopods will be crucial to determine whether these hardened structures are derived from conserved ancestral cell types or represent independent evolutionary acquisitions. Another related question concerns whether the ancestral mollusc possessed chaetae, as implied by the Cambrian fossil *Pelagiella exigua*, which potentially exhibited both chaetae and a mineralized shell (Thomas et al. 2020). Therefore, our findings indicate that extant molluscs retained components of the ancestral genetic architecture required for chaetal formation, which may have been secondarily co-opted, modified or lost during molluscan evolution. Together, these considerations highlight the need for integrative cellular comparisons across spiralian phyla to reconstruct the ancestral toolkit from which diverse hardened structures evolved.

In summary, our study provides a cellular and evolutionary framework for understanding the interplay between ancestral epithelial-secretory gene expression programs and lineage-specific effector diversification underlies the evolution of molluscan shells. By tracing the cellular and molecular connections between molluscan shells and other hardened structures across Spiralia, our findings highlight a general principle that evolutionary novelties emerge from the combinations, redeployments, and modifications of conserved cellular programs, with the incorporation of lineage-specific genes and duplicated paralogs. Moreover, derived cellular functions can evolve through extensive remodeling of pre-existing cellular systems without requiring the *de novo* origin of entirely novel cellular substrates. These advances provide insights into reconstructing the ancestral spiralian cell-type repertoire, revealing hidden ancestral traits, and distinguishing which morphological features were inherited from the LCA of animals and which evolved multiple times independently.

## Materials and methods

### scRNA-seq library preparation and sequencing

The mantle tissue was collected from an adult oyster obtained from the oyster farm in Laizhou (Shandong Province, China) and washed three times with ice-cold 1×PBS at 4 °C. The cleaned tissue was cut into ∼ 0.5 mm² fragments, washed again with 1×PBS, and digested on a shaker at 37 °C until complete dissociation. The suspension was filtered through a 40 µm strainer, and centrifuged at 300 g for 5 min at 4 °C. After removing the supernatant, cells were washed, centrifuged and resuspended with ice-cold 1×PBS. Cell concentration and viability were assessed using a cell counter, and the suspension was adjusted to 700-1200 cells/µL with >85% viability and <15% aggregation.

A single-cell RNA-seq library was generated using the BMKMANU DG1000 platform (Biomarker Technologies, China), a droplet-based single-cell RNA-sequencing system that uses microfluidic encapsulation of individual cells together with barcoded beads. The cell suspension was resuspended in PBS containing 0.04% BSA and loaded onto a BMKMANU C1000 chip using the DG1000 GEM Kit according to the manufacturer’s protocol. Individual cells and barcoded beads were co-encapsulated into gel bead-in-emulsion droplets (GEMs), in which cells were lysed and released transcripts were barcoded during reverse transcription. Barcoded cDNA was subsequently amplified and used for sequencing-library construction. Library concentration and quality were assessed using a Qubit 4.0 fluorometer (Thermo Fisher Scientific) and an Agilent 2100 (Agilent Technologies). One library was generated from the adult mantle sample and sequenced in a single sequencing run on the Illumina NovaSeq X Plus platform using 150-bp paired-end sequencing.

### Processing of scRNA-seq data

The BMKMANU sequencing output follows a barcode/UMI read structure that is compatible with the 10x Genomics Cell Ranger workflow. Accordingly, sequencing reads were subsequently processed using Cell Ranger. Raw sequencing data from the adult mantle tissue were processed with dealFQ v.1.3 (http://www.biomarker.com.cn/zhizao/tools) and then aligned to the *C. gigas* genome (GCF_902806645.1) (Penaloza et al. 2021) using CellRanger v.9.0.1 (https://github.com/10XGenomics/cellranger). The raw count matrix was generated and imported into Seurat v.5.3.0 (Stuart et al. 2019) for downstream analysis. Genes expressed in fewer than three cells and cells expressing fewer than 300 genes were removed. Low-quality cells were further filtered based on feature counts, total UMI counts, and mitochondrial gene percentage using median ± 3×MAD (median absolute deviation) thresholds. Filtered data were normalized using SCTransform (Choudhary and Satija 2022), and cell-cycle effects were regressed out using custom S-phase and G2/M-phase gene sets, identified by best-reciprocal hits with MMseqs2 v.17.b804f (Steinegger and Söding 2017) between *C. gigas* and *Homo sapiens*. Dimensionality reduction was performed using PCA, followed by UMAP visualization based on the K-nearest-neighbor graph (Hao et al. 2024). Clustering was performed across multiple resolutions, an optimal resolution of 1.2 was selected based on Clustree analysis (Zappia and Oshlack 2018). Doublets were identified and removed using DoubletFinder (McGinnis et al. 2019).

Following clustering, marker genes for each cell cluster were determined using the FindAllMarkers function in Seurat. Genes were considered markers if they showed positive differential expression, a log2 fold-change > 0.25, were expressed in at least 25% of cells in either comparison group, and had a *P* value < 0.05. As an additional post-clustering quality-control step, clusters 31 and 34 were excluded because they exhibited mixed transcriptional signatures inconsistent with coherent cell-type identities and showed relatively high transcript complexity and UMI counts (Supplementary file 11A-B). Specifically, cluster 31 co-expressed the hemocyte-associated marker *TCGP* and the myocyte marker *Hss35*, whereas cluster 34 co-expressed *TCGP* together with multiple cilia-associated markers, including *Tekt1*, *Crocc*, *Rsh4a-2*, and *Unc-13033* (Supplementary file 2). Consistent with these mixed lineage signatures, both clusters showed elevated distributions of nFeature_RNA and nCount_RNA (Supplementary file 11C-D). We therefore considered them residual multiplet-like populations that remained after cell-level doublet filtering and excluded them from subsequent cell-type annotation and comparative analyses. The remaining clusters were annotated using an evidence-based marker strategy that integrated marker gene expression with previously reported spatial, molecular, and functional information from oyster studies (Song et al. 2016; Li et al. 2021a, b, c; Zhu et al. 2021; Yu et al. 2022; Bai et al. 2023; Piovani et al. 2023; De La Forest Divonne et al. 2025; Ren et al. 2026) (Supplementary file 2). The R package Monocle3 v.1.4.26 (Cao et al. 2019) was used for pseudotime analysis to construct single-cell trajectories. Proliferative cells were specified as the root to orient pseudotime. Root specification was used solely to orient the pseudotime trajectory and does not, by itself, establish a lineage relationship between proliferative cells and SECs.

Single-cell transcriptomes of *C. gigas* trochophore were obtained from published datasets (NCBI Bioproject accession PRJNA967338). Quality control, normalization, dimension reduction, clustering, and marker gene identification were performed as described above, with an optimal resolution of 2.0. Cell types were annotated using an evidence-based strategy integrating marker gene expression with previously reported spatial, molecular, and functional information from oyster studies (Huan et al. 2013; Li et al. 2021a, b, c; Zhu et al. 2021; Piovani et al. 2023) (Supplementary file 2).

To assess transcriptional relationships between trochophore and adult mantle cell populations, we employed multiple complementary approaches, including Pearson and Spearman correlation of average gene expression, Jensen-Shannon divergence (JSD) to quantify overall expression distribution similarity, Jaccard indices to evaluate overlap of marker genes, and hypergeometric tests to determine statistical significance of shared marker gene sets. These complementary metrics were used to evaluate the correspondence between trochophore shell gland populations and adult SEC populations, with other broadly conserved cell types serving as internal references.

### Fluorescent *in situ* hybridization

FISH was performed to validate the spatial distributions of representative marker genes for SEC populations in the adult mantle of *C. gigas*, as previously described (Hu et al. 2025). Mantle tissues were sampled and fixed in 4% paraformaldehyde (PFA) at 4 °C, dehydrated through a graded ethanol series, cleared in xylene, then embedded in paraffin and sectioned at 10 μm thickness. Primers used for probe synthesis were designed using Primer Premier v.6.0 (Premier Biosoft International, Palo Alto, CA, USA) to amplify cDNA fragments of each marker gene (Supplementary file 12). Biotin-labeled antisense RNA probes for *rbp4b*, *Unc-25054*, and *Unc-42230*, and digoxigenin (DIG)-labeled antisense RNA probes for *BMPER*, *Unc-34425*, and *Unc-46765* were transcribed *in vitro* from the corresponding PCR-amplified fragments using the Roche SP6/T7 Transcription Kit (Roche, Germany). Sections were deparaffinized, hydrated, and then treated with 4 μg/mL proteinase K (Roche, Germany), and refixed with 4% PFA. Subsequently, the sections were acetylated with acetic anhydride and neutralized with triethanolamine for 10 min. Following pre-hybridization, sections were hybridized with the corresponding RNA probes overnight at 65 °C. After post-hybridization washes, sections were blocked with 1% blocking reagent (Roche, Germany). Sections hybridized with DIG-labeled probes were incubated with Alexa Fluor 488-conjugated anti-digoxigenin antibody (Jackson ImmunoResearch, USA), whereas those hybridized with biotin-labeled probes were incubated with Alexa Fluor 647-conjugated anti-biotin antibody (Jackson ImmunoResearch, USA). Antibodies were diluted 1:1,000 in blocking reagent and incubated overnight at 4 °C. Nuclei were counterstained with DAPI according to the manufacturer’s instructions. All the images were acquired with using a Leica TCS SP8 laser confocal microscope (Germany).

### Expression profiling of SMP genes

Larval and adult SMPs of *C. gigas* were identified by best-hit BLAST searches against the *C. gigas* genome database, using the previously reported sets of *C. gigas* SMPs (Zhao et al. 2018). Bulk RNA-seq datasets encompassing 37 developmental stages and 11 major tissues/organs of *C. gigas* were retrieved from published studies (Zhang et al. 2012; Lian et al. 2025). Raw reads were quality-filtered using fastp v.0.23.4 (Chen 2023) and gene expression (Transcripts per million, TPM) was quantified with salmon v.1.10.1 (Patro et al. 2017) using the reference transcript sequences. Expression matrices were normalized using the Trimmed Mean of M-values (TMM) method implemented in the Trinity utility scripts (Haas et al. 2013).

### Functional annotation and enrichment analysis

Protein-coding genes of *C. gigas* were represented by their longest isoforms, and were functionally annotated using emapper v.2.1.12 against the eggNOG 5.0 database (Huerta-Cepas et al. 2019). For functional comparisons between larval shell glands and adult SECs, marker genes were identified using the FindMarkers function in Seurat by comparing each cell type with all other cells. Genes were retained if they showed positive differential expression, a log2 fold-change > 0.25, expression in at least 25% of cells in either comparison group, and a *P* value < 0.05. GO enrichment analysis was perform using the R package clusterProfiler v.4.10.1 (Xu et al. 2024), against all expressed genes annotated by eggNOG database (Huerta-Cepas et al. 2019) as the background. GO terms with FDR-adjusted significance (*q* < 0.05) were considered statistically enriched.

### Phylostratigraphy analysis

Gene age was estimated using the GenERA v.1.4.2 (Barrera-Redondo et al. 2023), which implements genomic phylostratigraphy (Domazet-Lošo et al. 2007) by searching homologs in the NCBI non-redundant (NR) database (Release 2024_01_01) and mapping them onto the NCBI Taxonomy to assign an evolutionary origin to each gene. To quantify evolutionary signatures in expression, we calculated the transcriptome age index (TAI) using the R package myTAI (Drost et al. 2018), based on log-transformed average expression per cluster obtained with the AverageExpression function of Seurat. Phylostrata enrichment in cell type-specific marker genes was assessed with a hypergeometric test, comparing the observed distribution across phylostrata with that of the globally expressed gene set. The TAI contribution of each phylostratum for larval stages and adult tissues was calculated as previously described (Wang et al. 2020).

Duplicated genes and the phylogenetic timing of gene duplication events in *C. gigas* were inferred using OrthoFinder v2.5.2 (Emms and Kelly 2019) based on orthogroups and gene trees constructed across 40 metazoan species (Supplementary file 13), following a previously described approach (Bai et al. 2026). Duplicated genes were intersected with cell-type-specific marker genes from the trochophore and adult mantle datasets. Enrichment of duplicated genes at each duplication node was assessed using hypergeometric tests, with genes detected in the corresponding scRNA-seq dataset and represented in the OrthoFinder analysis used as the background. *P* values were adjusted for multiple testing using the Benjamini-Hochberg (BH) method. Enrichment was summarized as log2 fold-enrichment, calculated from the observed relative to expected frequency of duplicated genes at each duplication node. Functional enrichment of duplicated marker genes was further assessed by GO enrichment analysis as described above.

### Cross-species cell-type comparison

We re-analyzed the single-cell transcriptomes of the annelid worm *P. leidyi* (Alvarez-Campos et al. 2024) and the chaetognath *P. gotoi* (Piovani et al. 2025) using the same processing pipeline applied to the oyster mantle dataset. Pre-processed count matrices from the published study were retrieved for the flatworms *D. japonica* and *S. mediterranea* (Garcia-Castro et al. 2021). Seurat objects were generated according to the original cell annotation.

SAMap v.1.0.15 (Tarashansky et al. 2021) was applied between the *C. gigas* scRNA-seq dataset with datasets from *P. leidyi* (Alvarez-Campos et al. 2024), *D. japonica* (Garcia-Castro et al. 2021), *S. mediterranea* (Garcia-Castro et al. 2021), and *P. gotoi* (Piovani et al. 2025). Count matrices exported from Seurat in h5ad format were loaded into a SAMap object, and a BLAST-based homology graph was generated. Alignment scores between cell types were computed using the get_mapping_scores function, and homologous gene pairs were identified with GenePairFinder using a *P*-value of 0.05.

## Acknowledgements

This work was supported by National Natural Science Foundation of China (W2411019), Blue Seed Industry Innovation Project of Qingdao Institute of Blue Seed Industry (QDLYY-2024001), and Shandong Province (2025LZGC036).

## Author contributions

Yitian Bai, Conceptualization, Investigation, Methodology, Formal analysis, Writing – original draft; Biyang Hu, Investigation (FISH experiments); Kunyin Jiang, Sample collection, Data generation; Hong Yu, Resources, Methodology; Lingfeng Kong, Resources, Methodology; Shaojun Du, Funding acquisition, Writing – review and editing; Shikai Liu, Conceptualization, Supervision, Resources, Writing – review and editing; Qi Li, Conceptualization, Supervision, Funding acquisition, Resources, Writing – review and editing.

## Data availability

The scRNA-seq data generated in the current study have been deposited in the NCBI database under BioProject accession number PRJNA1338802. All original code has been deposited on GitHub at https://github.com/StevenBai97/Shell-cell-type-innovations.

---

Figure 1-figure supplement 1. Dual-color FISH reveals spatial segregation of transcriptionally related SEC populations in the adult oyster mantle. (A) Dual-color FISH for the SEC1 marker *rbp4b* (red) and the SEC2 marker *BMPER* (green). *rbp4b* expression is enriched along the pallial outer epithelium, whereas *BMPER* is concentrated in the outer-fold epithelium. Red and green arrowheads indicate representative *rbp4b* and *BMPER* expression domains, respectively. Nuclei are stained with DAPI (blue). Scale bar, 250 μm. (B) Dual-color FISH for the SEC4 marker *Unc-25054* (red) and the SEC3 marker *Unc-34425* (green). *Unc-34425* is predominantly expressed in the outer-fold epithelium, whereas *Unc-25054* shows a broad distribution along the pallial epithelium and the inner region of the outer fold. Red and green arrowheads indicate representative *Unc-25054* and *Unc-34425* expression domains, respectively, and yellow arrowheads indicate regions of spatial proximity or partial overlap. Nuclei are stained with DAPI (blue).

Figure 2-figure supplement 1. Dot plot showing the expression of representative marker genes across cell clusters of the *C. gigas* trochophore. Detailed information of marker genes used for cell-type annotation is provided in Supplementary file 2. Clusters lacking sufficiently characterized marker genes are classified as “unknown”.

Figure 2-figure supplement 2. Distinct expression profiles of larval and adult SMP genes across development and shell-forming cell populations of *C. gigas*. (A, D) Heatmaps showing the expression dynamics of larval (A) and adult (D) SMP genes across developmental stages and adult tissues. (B, E) Dot plots showing the expression of larval (B) and adult (E) SMP genes in each cell type from the trochophore scRNA-seq data. Shell-forming cell types are highlighted with gray backgrounds. (C, F) Dot plots showing the expression of larval (C) and adult (F) SMP genes in each cell type from scRNA-seq data of the adult mantle. Shell-forming cell types are highlighted with gray backgrounds. Abbreviations: TC, two-cell; FC, four-cell; EM, early morula; M, morula; B, blastula; RM, rotary movement; FS, free swimming; EG, early gastrula; G, gastrula; T1-T5, trochophore stages; ED1-ED2, early D-shape larva stages; D1-D7, D-shape larva stages; EU1-EU2, early umbo stages; U1-U6, umbo stages; LU1-LU2, late umbo stages; P1-P2, pediveliger stages; S, spat; J, juvenile; ME, mantle edge; MC, mantle center; SH, shell; He, hemolymph; AM, adductor muscle; Gi, gill; LP, labial palp; DG, digestive gland; MG, male gonad; FG, female gonad; Re, remaining tissues; SEC: Shell-secreting epithelial cells.

Figure 2-figure supplement 3. Pairwise transcriptomic comparisons of cell types between larval (trochophore) and adult (mantle) based on Pearson correlation (upper left), Spearman correlation (upper right), Jensen-Shannon similarity (1-JSD) (lower left), and Jaccard index (lower right). Shell glands and SECs are marked in red. Red dashed boxes indicate the comparisons between larval and adult shell-forming cell types. Abbreviations: SEC, shell-secreting epithelial cells.

Figure 3-figure supplement 1. Transcriptome Age Index (TAI) across oyster ontogeny (A) and adult tissues (B), computed using published bulk transcriptomes. Shaded areas indicate ± standard error. Abbreviations are as in Figure 2-figure supplement 2.

Figure 4-figure supplement 1. Re-analyses of cell atlases of two spiralian species. (A, C) Two-dimensional uniform manifold approximation and projection (UMAP) visualizations of cell clusters in the annelid worm *P. leidyi* (A) and the chaetognath *P. gotoi* (C). (B, D) Dot plots showing the expression of selected cell-type marker genes for each cluster in *P. leidyi* (B) and *P. gotoi* (D), according to previous studies (Alvarez-Campos et al. 2024; Piovani et al. 2025).

Figure 4-figure supplement 2. Conservation and functional enrichment of genes underlying cross-species transcriptional similarities of adult SECs in *C. gigas*. (A) Venn diagram showing the overlap of *C. gigas* genes involved in cross-species gene pairs linking SECs to transcriptionally similar cell types in *P. leidyi*, *D. japonica*, *S. mediterranea*, and *P. gotoi*. Gene pairs and their associated cell-type identities are provided in Supplementary file 9. Numbers indicate the numbers of *C. gigas* genes unique to or shared among the four cross-species comparisons. (B) GO enrichment analysis of the *C. gigas* genes involved in these cross-species gene pairs linking SECs to transcriptionally similar cell types in *P. leidyi*, *D. japonica*, *S. mediterranea*, and *P. gotoi*. Representative significantly enriched GO terms are grouped into major functional categories, including stress response, vesicle/lysosome-associated functions, translation/ribosome, ER protein targeting, and RNA metabolism. Dot size represents the gene ratio, and color indicates −log10(*q*-value). For visualization, −log10(*q*-value) values greater than 10 were capped at 10.

## Additional files

Supplementary file 1. The proportion of cell clusters in adult mantle of *C. gigas*.

Supplementary file 2. Marker genes for cell-type annotation in larval (trochophore) and adult (mantle) stages of *C. gigas*. Gene names used in this study were retained from previously published oyster studies where established. For genes requiring cross-species orthology assessment, reciprocal best BLAST hits were identified between the predicted proteomes of *C. gigas* and the corresponding reference species. Genes without sufficiently resolved functional annotations were retained using uncharacterized-gene identifiers.

Supplementary file 3. The proportion of cell clusters of *C. gigas* trochophore.

Supplementary file 4. Pairwise counts of overlapping marker gene betweem larval (trochophore) and adult (mantle) cell types. Values indicate the numbers of overlapping marker genes between each pair of cell types. Numbers in parentheses are statistical significance of marker-gene overlap assessed using a one-sided hypergeometric test. Red shading indicates significant overlaps (BH-adjusted *P* < 0.05).

Supplementary file 5. GO enrichment of marker genes in larval shell gland and adult SEC of *C. gigas* (FDR-adjusted significance, *q* value < 0.05). The GO term shared between SECs and shell gland cells is highlighted in red.

Supplementary file 6. Protein-domain annotations of marker genes in larval shell gland and adult SEC of *C. gigas*.

Supplementary file 7. Gene duplication among marker genes of larval and adult shell-forming cell populations in *C. gigas*. Genes associated with inferred duplication events are listed together with their assigned phylogenetic duplication nodes. A gene may be assigned to multiple duplication nodes when its evolutionary history includes multiple inferred duplication events.

Supplementary file 8. GO enrichment of duplicated marker genes in larval shell gland and adult SEC of *C. gigas* (FDR-adjusted significance, *q* value < 0.05).

Supplementary file 9. Cross-species gene pairs supporting cell-population transcriptional similarities between *C. gigas* and four other spiralian species.

Supplementary file 10. Cross-species gene pairs supporting transcriptional similarities between merged cell types of *C. gigas* and four other spiralian species. For the SAMAP analysis, function related cell types were merged into broader cell-type categories prior to cross-species comparison. For example, the six SEC populations were merged as “SEC”.

Supplementary file 11. Post-clustering quality control of the adult mantle single-cell atlas in *C. gigas*.

(A) UMAP visualization of the initial 35 transcriptional clusters identified in the adult mantle before post-clustering quality control.

(B) Dot plot showing the expression of representative marker genes across the initial clusters. Red arrowheads indicate clusters 31 and 34, which exhibited mixed expression of markers associated with otherwise distinct cell lineages. In particular, cluster 31 co-expressed the hemocyte-associated marker *TCGP* and the myocyte marker *Hss35*, whereas cluster 34 co-expressed *TCGP* together with multiple cilia-associated markers, including *Tekt1*, *Crocc*, *Rsh4a-2*, and *Unc-13033*.

(C, D) Violin plots showing the distributions of the number of detected genes per cell (nFeature_RNA; C) and total UMI counts per cell (nCount_RNA; D) across clusters. Clusters 31 and 34 are highlighted in red and showed relatively elevated transcript complexity and UMI counts compared with most other clusters.

Supplementary file 12. Primers for FISH used in this study. T7 promoter sequences were underlined.

Supplementary file 13. Genome resources used for the identification of duplicated genes by OrthoFinder analysis.

## Notes

### Competing Interest Statement

The authors have declared no competing interest.

### Summary of Updates

This version has been substantially revised in response to peer review. We performed fluorescence in situ hybridization experiments to validate the spatial distributions of six shell secreting epithelial cell populations. We re-analyzed the trochophore single-cell dataset and expanded the larval and adult comparison to include protein domain analysis. Cross-species comparisons were revised to examine homologous gene pairs and their functional enrichment. We also added analyses of lineage-specific gene duplication and expanded the phylostratigraphic analysis. The evolutionary interpretations have been refined to distinguish transcriptional similarity, functional conservation, and cell type homology. Figures, supplementary files, and methodological descriptions have been updated accordingly. Biyang Hu has been added as a co-author in recognition of her contribution to the fluorescence in situ hybridization experiments. All authors have agreed to this authorship change.

## Reference

Aguilera F, McDougall C, Degnan BM. 2017. Co-Option and De Novo Gene Evolution Underlie Molluscan Shell Diversity. Molecular Biology and Evolution 34: 779–792.

Alvarez-Campos P, Garcia-Castro H, Emili E, Perez-Posada A, Del Olmo I, Peron S, Salamanca-Diaz DA, Mason V, Metzger B, Bely AE et al. 2024. Annelid adult cell type diversity and their pluripotent cellular origins. Nature Communications 15: 3194.

Arendt D, Musser JM, Baker CVH, Bergman A, Cepko C, Erwin DH, Pavlicev M, Schlosser G, Widder S, Laubichler MD et al. 2016. The origin and evolution of cell types. Nature Reviews Genetics 17: 744–757.

Arivalagan J, Yarra T, Marie B, Sleight VA, Duvernois-Berthet E, Clark MS, Marie A, Berland S. 2017. Insights from the Shell Proteome: Biomineralization to Adaptation. Molecular Biology and Evolution 34: 66–77.

Bai Y, Liu S, Hu Y, Yu H, Kong L, Xu C, Li Q. 2023. Multi-omic insights into the formation and evolution of a novel shell microstructure in oysters. BMC Biology 21: 204.

Bai Y, Min Y, Liu S, Hu Y, Jin S, Yu H, Kong L, Macqueen DJ, Du S, Li Q et al. 2026. Evolutionary innovation within conserved gene regulatory networks underlying biomineralized skeletons in Bilateria. Molecular Biology and Evolution 43: 1–23.

Barrera-Redondo J, Lotharukpong JS, Drost H-G, Coelho SM. 2023. Uncovering gene-family founder events during major evolutionary transitions in animals, plants and fungi using GenEra. Genome Biology 24: 54.

Barrera Grijalba CC, Rodriguez Monje SV, Ariza Aranguren G, Lunzer K, Scherholz M, Redl E, Wollesen T. 2025. Molluscan shells, spicules, and gladii are evolutionarily deeply conserved. Journal of Experimental Zoology Part B: Molecular and Developmental Evolution 344: 198–213.

Cao J, Spielmann M, Qiu X, Huang X, Ibrahim DM, Hill AJ, Zhang F, Mundlos S, Christiansen L, Steemers FJ et al. 2019. The single-cell transcriptional landscape of mammalian organogenesis. Nature 566: 496–502.

Carscadden KA, Batstone RT, Hauser FE. 2023. Origins and evolution of biological novelty. Biological Reviews 98: 1472–1491.

Cavallo A, Clark MS, Peck LS, Harper EM, Sleight VA. 2022. Evolutionary conservation and divergence of the transcriptional regulation of bivalve shell secretion across life-history stages. Royal Society Open Science 9: 221022.

Chen S. 2023.Ultrafast one-pass FASTQ data preprocessing, quality control, and deduplication using fastp. iMeta 2: e107.

Chen Z, Baeza JA, Chen C, Gonzalez MT, González VL, Greve C, Kocot KM, Arbizu PM, Moles J, Schell T et al. 2025. A genome-based phylogeny for Mollusca is concordant with fossils and morphology. Science 387: 1001–1007.

Choudhary S, Satija R. 2022. Comparison and evaluation of statistical error models for scRNA-seq. Genome Biology 23: 27.

Clark MS, Peck LS, Arivalagan J, Backeljau T, Berland S, Cardoso JCR, Caurcel C, Chapelle G, De Noia M, Dupont S et al. 2020. Deciphering mollusc shell production: the roles of genetic mechanisms through to ecology, aquaculture and biomimetics. Biological Reviews 95: 1812–1837.

De La Forest Divonne S, Pouzadoux J, Romatif O, Montagnani C, Mitta G, Destoumieux-Garzón D, Gourbal B, Charriere GM, Vignal E. 2025. Diversity and functional specialization of oyster immune cells uncovered by integrative single-cell level investigations. eLife 13: RP102622.

Domazet-Lošo T, Brajković J, Tautz D. 2007. A phylostratigraphy approach to uncover the genomic history of major adaptations in metazoan lineages. Trends in Genetics 23: 533–539.

Drost H-G, Gabel A, Liu J, Quint M, Grosse I. 2018. myTAI: evolutionary transcriptomics with R. Bioinformatics 34: 1589–1590.

Emms DM, Kelly S. 2019. OrthoFinder: phylogenetic orthology inference for comparative genomics. Genome Biology. 20: 238.

Erwin DH. 2021. A conceptual framework of evolutionary novelty and innovation. Biological Reviews 96: 1–15.

Garcia-Castro H, Kenny NJ, Iglesias M, Alvarez-Campos P, Mason V, Elek A, Schonauer A, Sleight VA, Neiro J, Aboobaker A et al. 2021. ACME dissociation: a versatile cell fixation-dissociation method for single-cell transcriptomics. Genome Biology 22: 89.

Haas BJ, Papanicolaou A, Yassour M, Grabherr M, Blood PD, Bowden J, Couger MB, Eccles D, Li B, Lieber M et al. 2013. De novo transcript sequence reconstruction from RNA-seq using the Trinity platform for reference generation and analysis. Nature Protocols 8: 1494–1512.

Hao Y, Stuart T, Kowalski MH, Choudhary S, Hoffman P, Hartman A, Srivastava A, Molla G, Madad S, Fernandez-Granda C et al. 2024. Dictionary learning for integrative, multimodal and scalable single-cell analysis. Nature Biotechnology 42: 293–304.

Herlitze I, Marie B, Marin F, Jackson DJ. 2018. Molecular modularity and asymmetry of the molluscan mantle revealed by a gene expression atlas. Gigascience 7: giy056.

Hu B, Yu H, Kong L, Liu S, Du S, Li Qi. 2025. *Cg*PPOX and *Cg*FECH mediate protoporphyrin ix accumulation predominantly in the outer mantle fold of Pacific oyster *Crassostrea gigas*. Marine Biotechnology 27: 46.

Huan P, Liu G, Wang H, Liu B. 2013. Identification of a tyrosinase gene potentially involved in early larval shell biogenesis of the Pacific oyster *Crassostrea gigas*. Development Genes And Evolution 223: 389–394.

Huerta-Cepas J, Szklarczyk D, Heller D, Hernández-Plaza A, Forslund SK, Cook H, Mende DR, Letunic I, Rattei T, Jensen Lars J et al. 2019. eggNOG 5.0: a hierarchical, functionally and phylogenetically annotated orthology resource based on 5090 organisms and 2502 viruses. Nucleic Acids Research 47: D309–D314.

Jackson DJ, McDougall C, Green K, Simpson F, Worheide G, Degnan BM. 2006. A rapidly evolving secretome builds and patterns a sea shell. BMC Biology 4: 40.

Jackson DJ, McDougall C, Woodcroft B, Moase P, Rose RA, Kube M, Reinhardt R, Rokhsar DS, Montagnani C, Joubert C et al. 2010. Parallel evolution of nacre building gene sets in molluscs. Molecular Biology and Evolution 27: 591–608.

Kocot KM, Aguilera F, McDougall C, Jackson DJ, Degnan BM. 2016. Sea shell diversity and rapidly evolving secretomes: insights into the evolution of biomineralization. Frontiers in Zoology 13: 23.

Korsunsky I, Millard N, Fan J, Slowikowski K, Zhang F, Wei K, Baglaenko Y, Brenner M, Loh P-r, Raychaudhuri S. 2019. Fast, sensitive and accurate integration of single-cell data with Harmony. Nature Methods 16: 1289–1296.

Laumer Christopher E, Bekkouche N, Kerbl A, Goetz F, Neves Ricardo C, Sørensen Martin V, Kristensen Reinhardt M, Hejnol A, Dunn Casey W, Giribet G et al. 2015. Spiralian phylogeny informs the evolution of microscopic lineages. Current Biology 25: 2000–2006.

Li H, Li Q, Yu H, Du S. 2021a. Characterization of paramyosin protein structure and gene expression during myogenesis in Pacific oyster (*Crassostrea gigas*). Comparative Biochemistry and Physiology Part B: Biochemistry and Molecular Biology 255: 110594.

Li H, Yu H, Du S, Li Q. 2021b. CRISPR/Cas9 mediated high efficiency knockout of myosin essential light chain gene in the Pacific oyster (*Crassostrea gigas*). Marine Biotechnology 23: 215–224.

Li H, Yu H, Li Q. 2021c. Striated myosin heavy chain gene is a crucial regulator of larval myogenesis in the Pacific oyster *Crassostrea gigas*. International Journal of Biological Macromolecules 179: 388–397.

Li Y, Hu M, Zhang Z, Wu B, Zheng J, Zhang F, Hao J, Xue T, Li Z, Zhu C et al. 2025. Origin and stepwise evolution of vertebrate lungs. Nat Ecol Evol 9: 672–691.

Li Z, Yang M, Zhou C, Shi P, Hu P, Liang B, Jiang Q, Zhang L, Liu X, Lai C et al. 2024. Deciphering the molecular toolkit: regulatory elements governing shell biomineralization in marine molluscs. Integrative Zoology 20: 448–464.

Lian S, Hu N, Chen X, Dai X, Zhu X, Qiao R, Liu S, Lu Y, Zhang F, Sun F et al. 2025. Widespread presence of bone marrow–like hematopoietic stem cell niche in invertebrate skeletons. Science Advances 11: eadw0958.

Liu G, Huan P, Liu B. 2020. Identification of three cell populations from the shell gland of a bivalve mollusc. Development Genes and Evolution 230: 39–45.

Luo YJ, Takeuchi T, Koyanagi R, Yamada L, Kanda M, Khalturina M, Fujie M, Yamasaki SI, Endo K, Satoh N. 2015. The Lingula genome provides insights into brachiopod evolution and the origin of phosphate biomineralization. Nature Communications 6: 8301.

Marie B, Joubert C, Tayale A, Zanella-Cleon I, Belliard C, Piquemal D, Cochennec-Laureau N, Marin F, Gueguen Y, Montagnani C. 2012. Different secretory repertoires control the biomineralization processes of prism and nacre deposition of the pearl oyster shell. Proceedings of the National Academy of Sciences 109: 20986–20991.

Martin-Zamora FM, Liang Y, Guynes K, Carrillo-Baltodano AM, Davies BE, Donnellan RD, Tan Y, Moggioli G, Seudre O, Tran M et al. 2023. Annelid functional genomics reveal the origins of bilaterian life cycles. Nature 615: 105–110.

McDougall C, Degnan BM. 2018. The evolution of mollusc shells. WIREs Developmental Biology 7: e313.

McGinnis CS, Murrow LM, Gartner ZJ. 2019. DoubletFinder: Doublet detection in single-cell rna sequencing data using artificial nearest neighbors. Cell Systems 8: 329–337.e324.

Moczek AP. 2008. On the origins of novelty in development and evolution. BioEssays 30: 432–447.

Murdock DJE. 2020. The’biomineralization toolkit’ and the origin of animal skeletons. Biological Reviews 95: 1372–1392.

Parey E, Ortega-Martinez O, Delroisse J, Piovani L, Czarkwiani A, Dylus D, Arya S, Dupont S, Thorndyke M, Larsson T et al. 2024. The brittle star genome illuminates the genetic basis of animal appendage regeneration. Nature Ecology & Evolution 8: 1505–1521.

Parker J, Pennell M. 2025. The cellular substrate of evolutionary novelty. Curr Biol 35: R626–R637.

Patro R, Duggal G, Love MI, Irizarry RA, Kingsford C. 2017. Salmon provides fast and bias-aware quantification of transcript expression. Nature Methods 14: 417–419.

Penaloza C, Gutierrez AP, Eory L, Wang S, Guo X, Archibald AL, Bean TP, Houston RD. 2021. A chromosome-level genome assembly for the Pacific oyster Crassostrea gigas. Gigascience 10: giab020.

Peterson KJ, Lyons JB, Nowak KS, Takacs CM, Wargo MJ, McPeek MA. 2004. Estimating metazoan divergence times with a molecular clock. Proceedings of the National Academy of Sciences 101: 6536–6541.

Piovani L, Gavriouchkina D, Parey E, Sarre LA, Peijnenburg KTCA, Martín-Durán JM, Rokhsar DS, Satoh N, de Mendoza A, Goto T et al. 2025. The genomic origin of the unique chaetognath body plan. Nature 645: 964–973.

Piovani L, Leite DJ, Yañez Guerra LA, Simpson F, Musser JM, Salvador-Martínez I, Marlétaz F, Jékely G, Telford MJ. 2023. Single-cell atlases of two lophotrochozoan larvae highlight their complex evolutionary histories. Science Advances 9: eadg6034.

Piovani L, Marlétaz F. 2023. Single-cell transcriptomics refuels the exploration of spiralian biology. Briefings in Functional Genomics 22: 517–524.

Ren L, Bai Y, Shi C, Hao Z, Li Q, Macqueen DJ, Liu S. 2026. Glycophagy is an ancient bilaterian pathway supporting metabolic adaptation through STBD1 structural evolution. Communications Biology 9: 268.

Shimizu K, Takeuchi T, Negishi L, Kurumizaka H, Kuriyama I, Endo K, Suzuki M. 2022. Evolution of epidermal growth factor (EGF)-like and zona pellucida domains containing shell matrix proteins in mollusks. Molecular Biology and Evolution 39: msac089.

Sleight VA. 2023. Cell type and gene regulatory network approaches in the evolution of spiralian biomineralisation. Brief Funct Genomics 22: 509–516.

Sleight VA, Antczak P, Falciani F, Clark MS. 2020. Computationally predicted gene regulatory networks in molluscan biomineralization identify extracellular matrix production and ion transportation pathways. Bioinformatics 36: 1326–1332.

Song X, Wang H, Chen H, Sun M, Liang Z, Wang L, Song L. 2016. Conserved hemopoietic transcription factor Cg-SCL delineates hematopoiesis of Pacific oyster *Crassostrea gigas*. Fish & Shellfish Immunology 51:180–188.

Steinegger M, Söding J. 2017. MMseqs2 enables sensitive protein sequence searching for the analysis of massive data sets. Nature Biotechnology 35: 1026–1028.

Stuart T, Butler A, Hoffman P, Hafemeister C, Papalexi E, Mauck WM, III, Hao Y, Stoeckius M, Smibert P, Satija R. 2019. Comprehensive Integration of Single-Cell Data. Cell 177: 1888–1902.e1821.

Sun J, Chen C, Miyamoto N, Li R, Sigwart JD, Xu T, Sun Y, Wong WC, Ip JCH, Zhang W et al. 2020. The Scaly-foot Snail genome and implications for the origins of biomineralised armour. Nature Communications 11: 1657.

Sun Y, Sun J, Yang Y, Lan Y, Ip JC, Wong WC, Kwan YH, Zhang Y, Han Z, Qiu JW et al. 2021. Genomic signatures supporting the symbiosis and formation of chitinous tube in the deep-sea tubeworm *Paraescarpia echinospica*. Molecular Biology and Evolution 38: 4116–4134.

Tanay A, Sebé-Pedrós A. 2021. Evolutionary cell type mapping with single-cell genomics. Trends in Genetics 37: 919–932.

Tarashansky AJ, Musser JM, Khariton M, Li P, Arendt D, Quake SR, Wang B. 2021. Mapping single-cell atlases throughout Metazoa unravels cell type evolution. eLife 10.

Thomas RDK, Runnegar B, Matt K, Sigwart J. 2020. Pelagiella exigua, an early Cambrian stem gastropod with chaetae: lophotrochozoan heritage and conchiferan novelty. Palaeontology 63: 601–627.

Tommasini D, Yoshimatsu T, Puthussery T, Baden T, Shekhar K. 2025. Comparative transcriptomic insights into the evolution of vertebrate photoreceptor types. Current Biology 35: 2228–2239.e2224.

Wagner GP, Lynch VJ. 2010. Evolutionary novelties. Current Biology 20: R48–R52.

Wang J, Zhang L, Lian S, Qin Z, Zhu X, Dai X, Huang Z, Ke C, Zhou Z, Wei J et al. 2020. Evolutionary transcriptomics of metazoan biphasic life cycle supports a single intercalation origin of metazoan larvae. Nature Ecology & Evolution 4: 725–736.

Wanninger A, Wollesen T. 2019. The evolution of molluscs. Biological Reviews 94: 102–115.

Wernstrom JV, Gasiorowski L, Hejnol A. 2022. Brachiopod and mollusc biomineralisation is a conserved process that was lost in the phoronid-bryozoan stem lineage. Evodevo 13: 17.

Wu L, Ferger KE, Lambert JD. 2019. Gene expression does not support the developmental hourglass model in three animals with spiralian development. Mol Biol Evol 36: 1373–1383.

Xia S, Chen J, Arsala D, Emerson JJ, Long M. 2025. Functional innovation through new genes as a general evolutionary process. Nature Genetics 57: 295–309.

Xiong L-L, Niu R-Z, Chen L, Huangfu L-R, Li J, Xue L-L, Sun Y-F, Wang L-M, Li Y-P, Liu J et al. 2025. Cross-species insights from single-nucleus sequencing highlight aging-related hippocampal features in tree shrew. Molecular Biology and Evolution 42: msaf020.

Xu S, Hu E, Cai Y, Xie Z, Luo X, Zhan L, Tang W, Wang Q, Liu B, Wang R et al. 2024. Using clusterProfiler to characterize multiomics data. Nature Protocols 19: 3292–3320.

Yarra T, Blaxter M, Clark MS. 2021. A Bivalve Biomineralization Toolbox. Molecular Biology and Evolution 38: 4043–4055.

Yu S, Qiao X, Song X, Yang Y, Zhang D, Sun W, Wang L, Song L. 2022. The proliferating cell nuclear antigen (*PCNA*) is a potential proliferative marker in oyster *Crassostrea gigas*. Fish & Shellfish Immunology 122: 306–315.

Zappia L, Oshlack A. 2018. Clustering trees: a visualization for evaluating clusterings at multiple resolutions. GigaScience 7: giy083.

Zhang G, Fang X, Guo X, Li L, Luo R, Xu F, Yang P, Zhang L, Wang X, Qi H et al. 2012. The oyster genome reveals stress adaptation and complexity of shell formation. Nature 490: 49–54.

Zhao R, Takeuchi T, Luo YJ, Ishikawa A, Kobayashi T, Koyanagi R, Villar-Briones A, Yamada L, Sawada H, Iwanaga S et al. 2018. Dual gene repertoires for larval and adult shells reveal molecules essential for molluscan shell formation. Molecular Biology and Evolution 35: 2751–2761.

Zhong H, Han W, Gomez-Cabrero D, Tegner J, Gao X, Cui G, Aranda M. 2025. Benchmarking cross-species single-cell RNA-seq data integration methods: towards a cell type tree of life. Nucleic Acids Research 53.

Zhu Y, Li Q, Yu H, Liu S, Kong L. 2021. Shell biosynthesis and pigmentation as revealed by the expression of tyrosinase and tyrosinase-like protein genes in Pacific oyster (*Crassostrea gigas*) with different shell colors. Marine Biotechnology 23: 777–789.

