## Supplementary figures and images for "Ancestral secretory programs underlie the evolution of morphological innovations across Spiralia"

### Figure 1-figure supplement 1.jpg

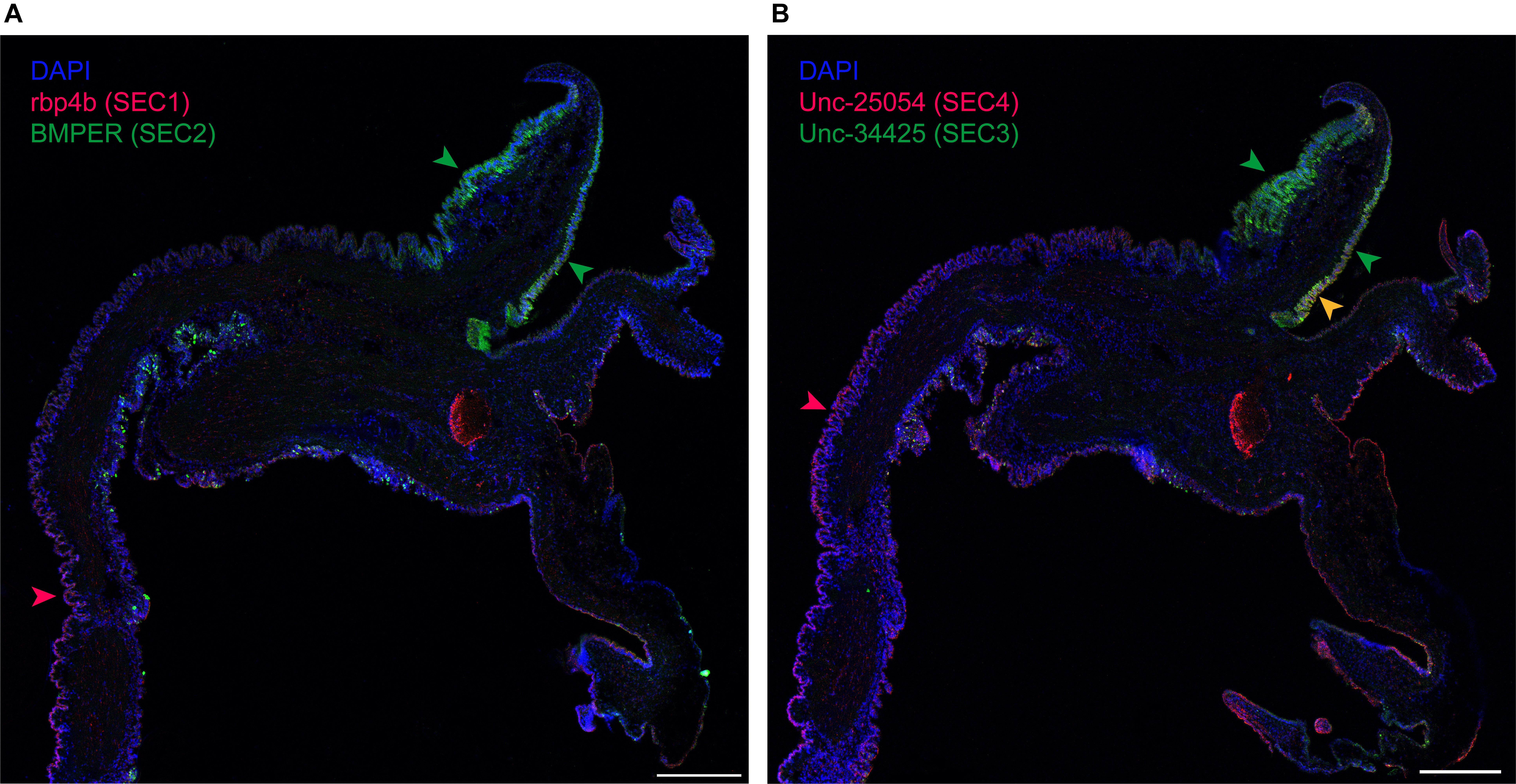

### Figure 2-figure supplement 1.jpg

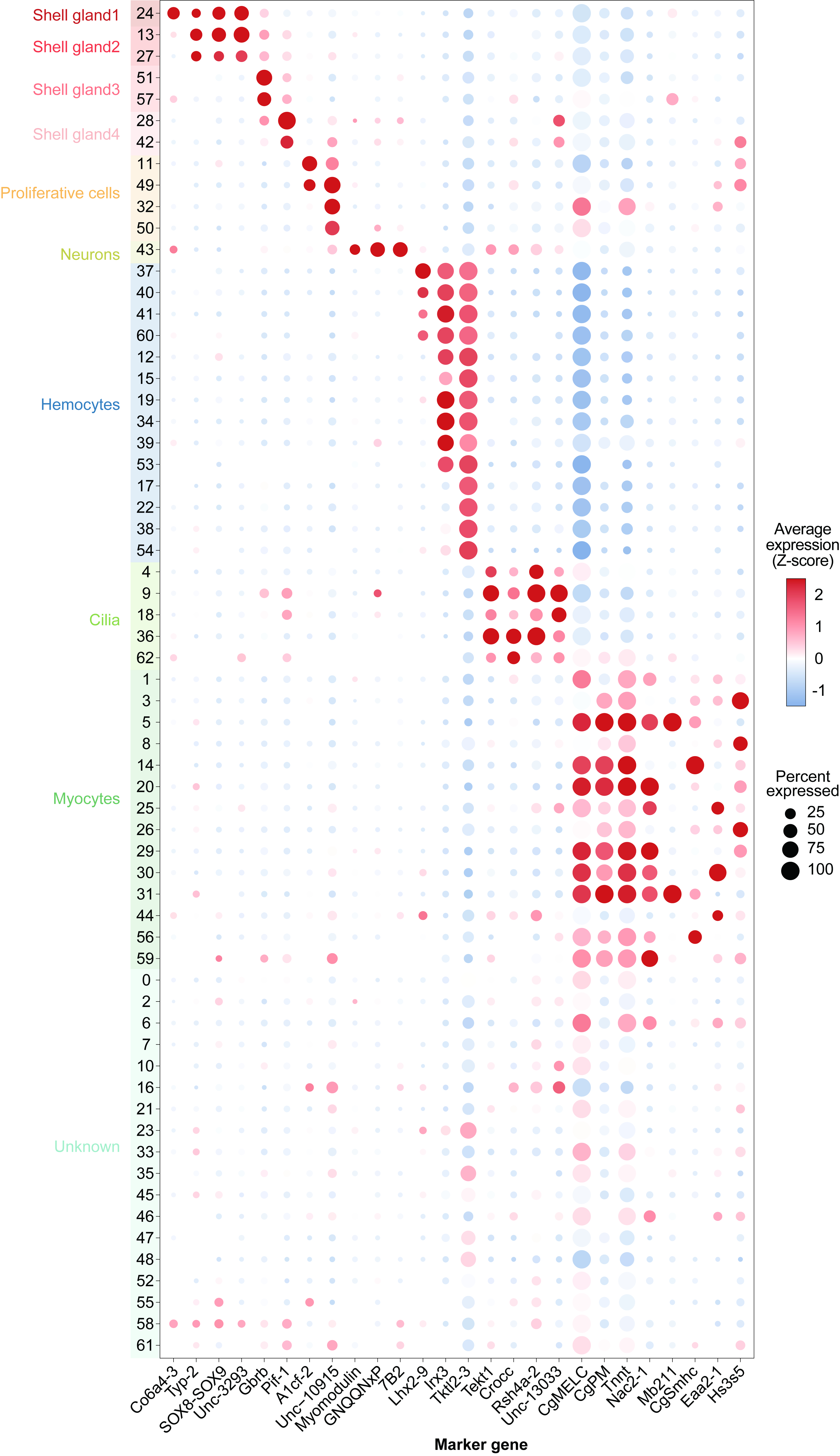

### Figure 2-figure supplement 2.jpg

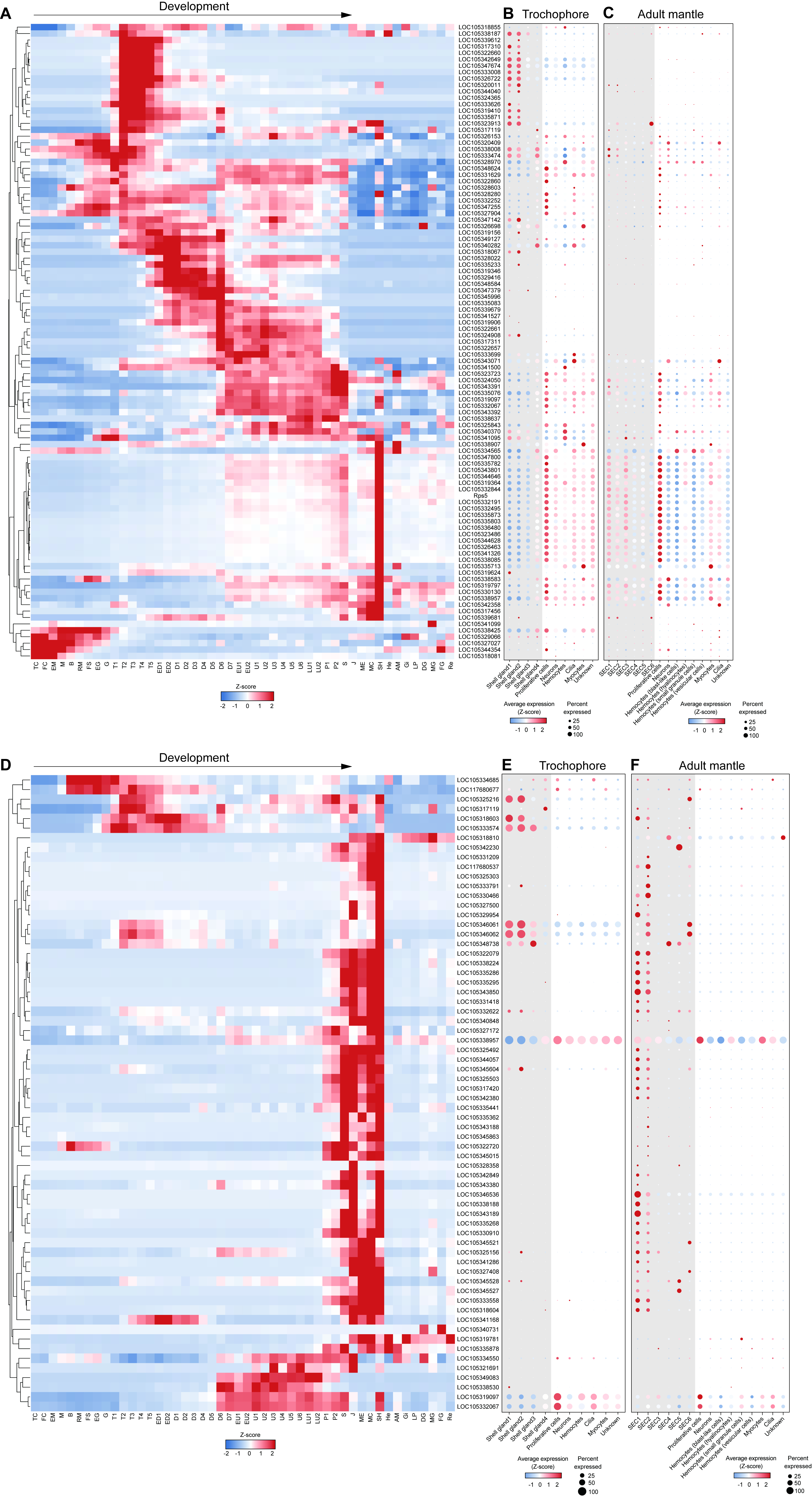

### Figure 2-figure supplement 3.jpg

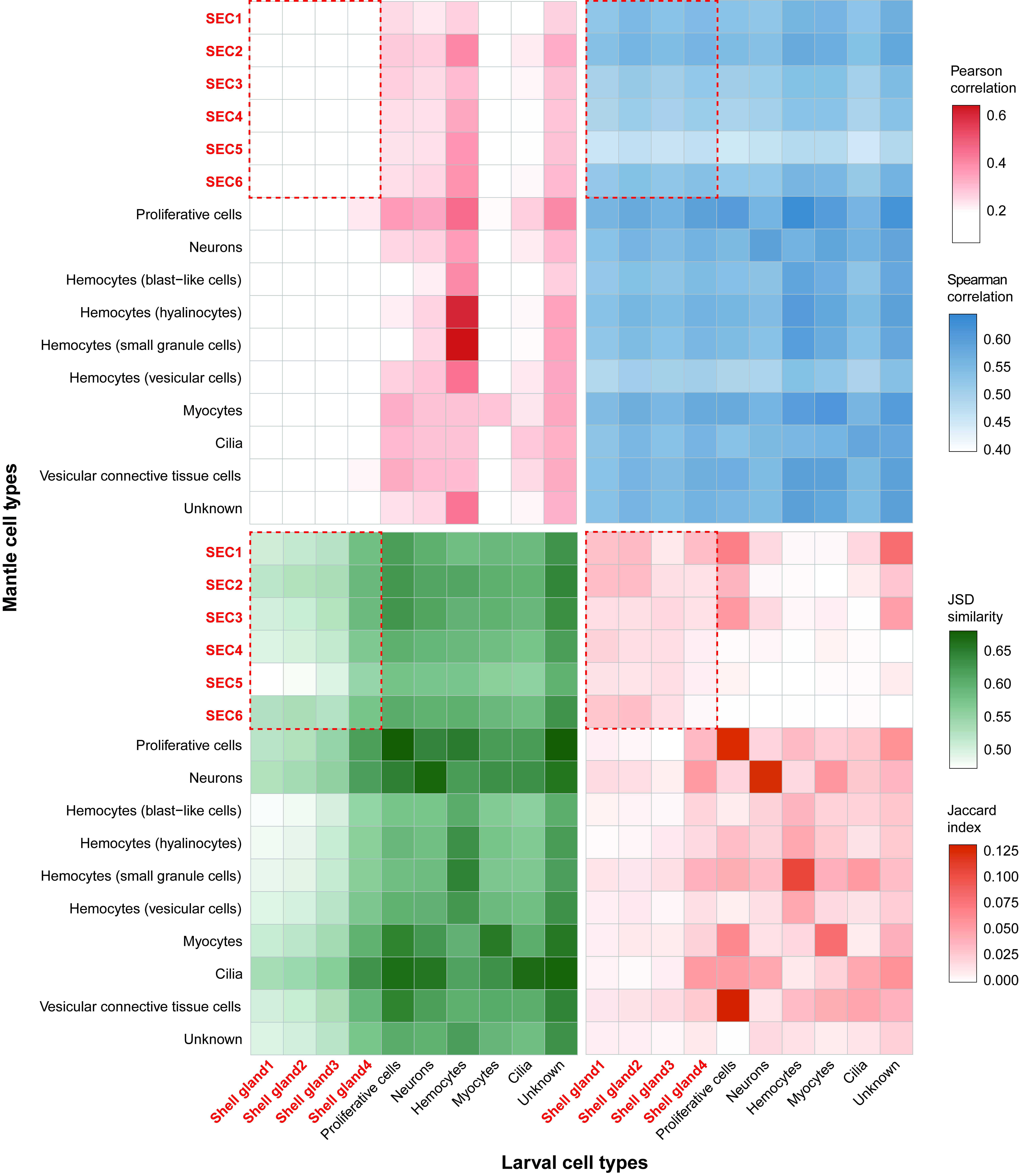

### Figure 3-figure supplement 1.jpg

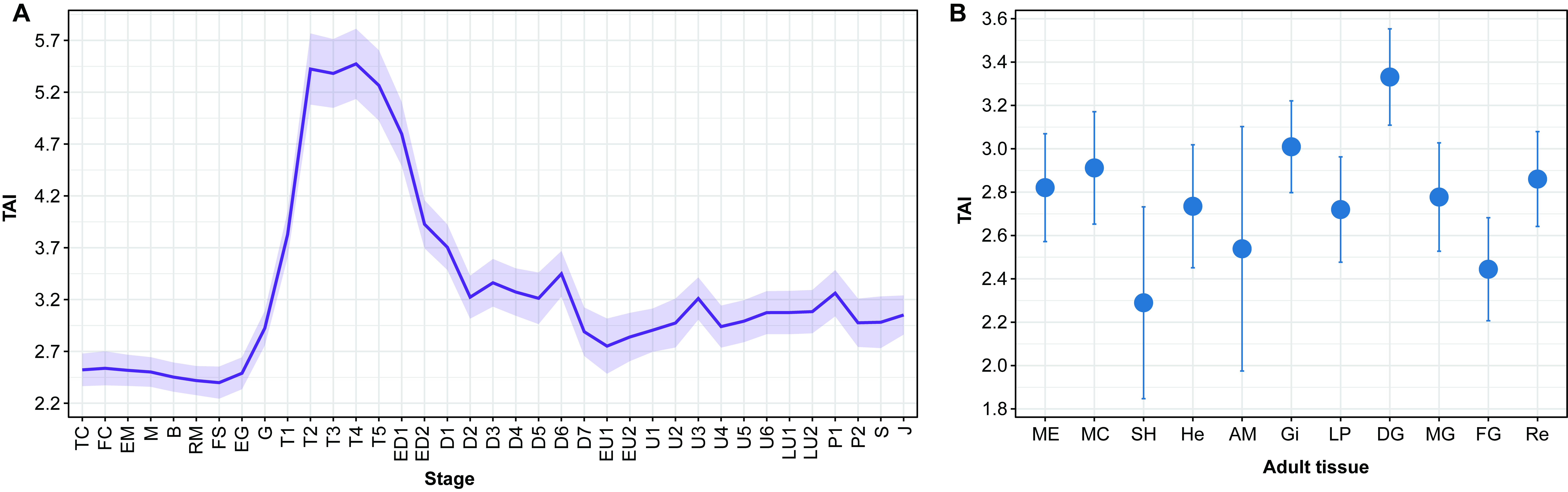

### Figure 4-figure supplement 1.jpg

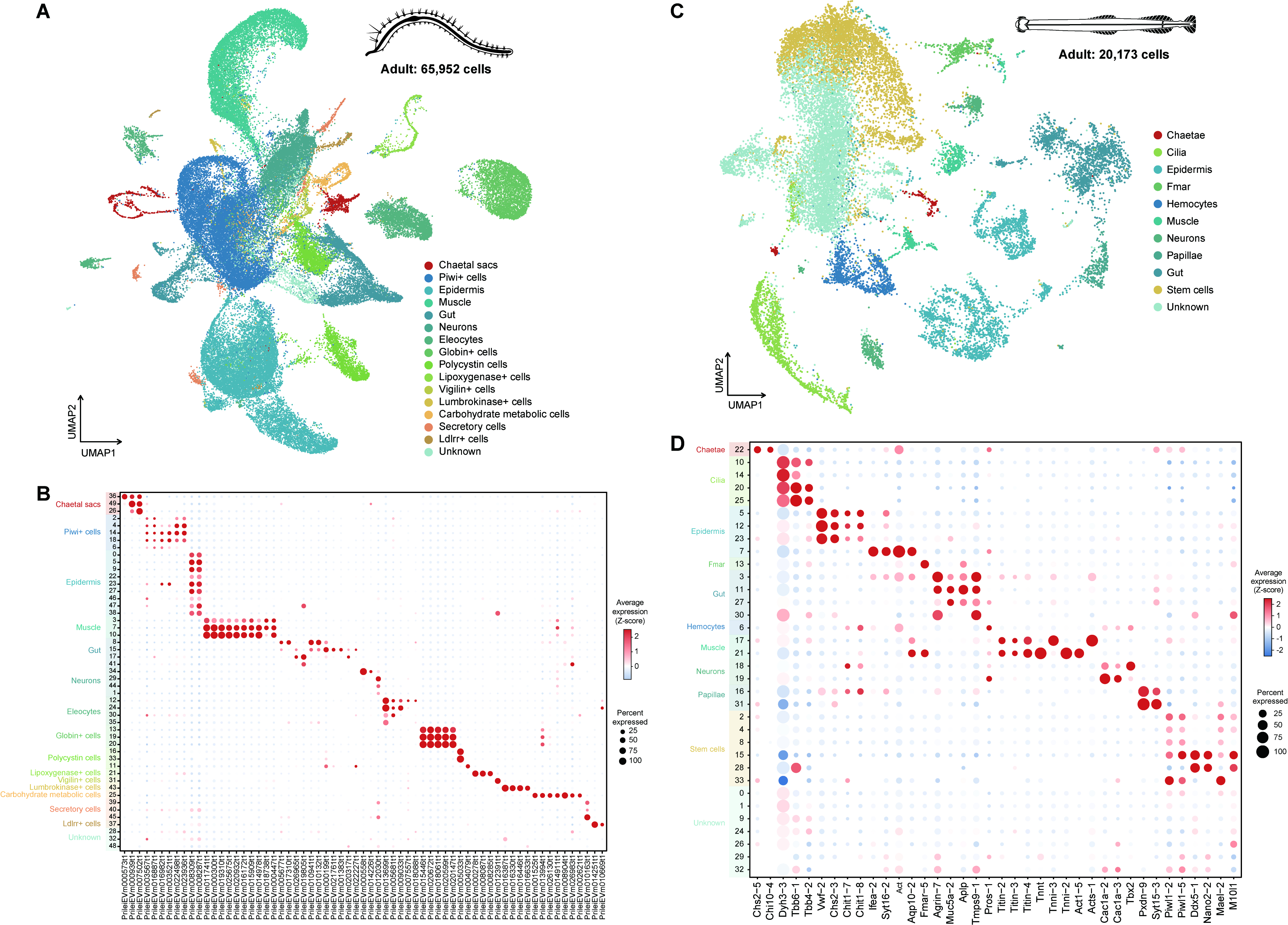

### Figure 4-figure supplement 2.jpg

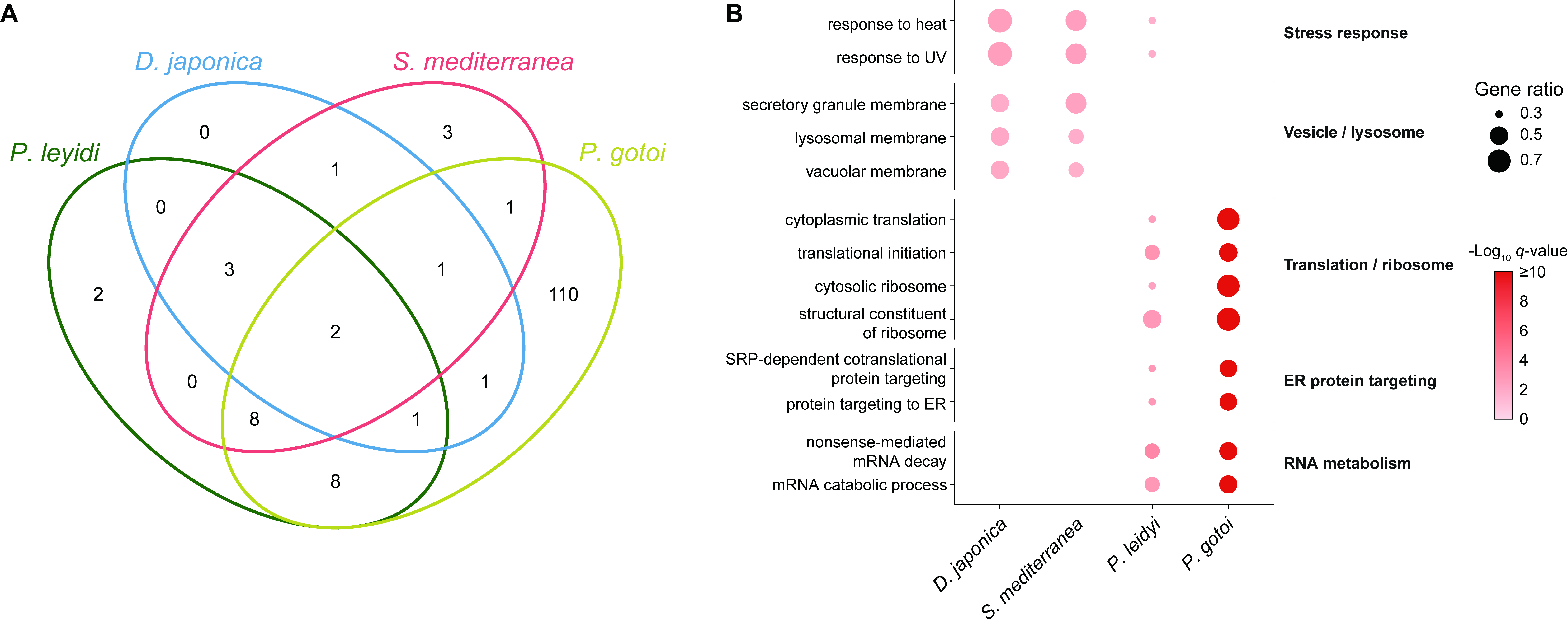

### Supplementary file 11.jpg

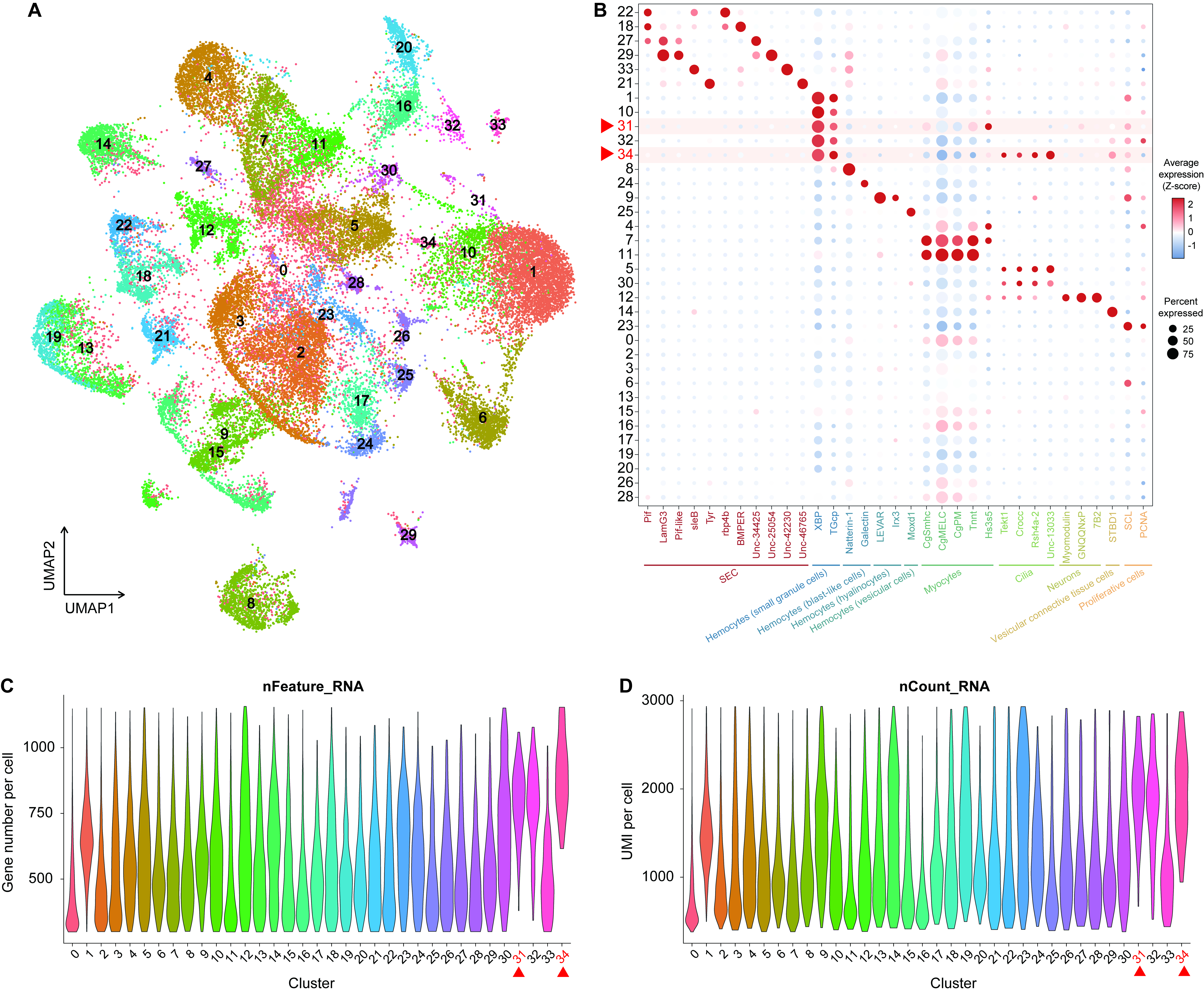
